# CHiMP: Deep Learning Tools Trained on Protein Crystallisation Micrographs to Enable Automation of Experiments

**DOI:** 10.1101/2024.05.22.595345

**Authors:** Oliver N. F. King, Karl E. Levik, James Sandy, Mark Basham

## Abstract

A group of three deep learning tools, referred to collectively as CHiMP (Crystal Hits in My Plate) were created for analysis of micrographs of protein crystallisation experiments at the Diamond Light Source (DLS) synchrotron, UK. The first tool, a classification network, assigns images into categories relating to experimental outcomes. The other two tools are networks that perform both object detection and instance segmentation, resulting in masks of individual crystals in the first case, and masks of crystallisation droplets in addition to crystals in the second case, allowing positions and sizes of these entities to be recorded. Creation of these tools used transfer learning, where weights from a pre-trained deep learning network were used as a starting point and re-purposed by further training on a relatively small set of data. Two of the tools are now integrated at the VMXi macromolecular crystallography beamline at DLS where they absolve the need for any user input both for monitoring crystallisation experiments and for triggering in situ data collections. The third is being integrated into the XChem fragment-based drug discovery screening platform, also at DLS, to allow automatic targeting of acoustic compound dispensing into crystallisation droplets.

## Background and Introduction

Determination of the three-dimensional structure of proteins by X-ray crystallography is a widely-adopted technique that provides invaluable information to allow the function of proteins to be elucidated. It is also capable of providing empirical evidence of the binding of ligands, such as enzyme substrates, co-factors or drug-like molecules that may modulate enzymatic activity. Despite the recent revolution in protein structure prediction by artificial intelligence (AI) such as Alphafold (1) and cryo electron microscopy (2), X-ray crystallography is still the technique capable of providing the highest resolution atomic coordinate information for these macromolecules, thereby enabling analysis of biochemical reactions and binding interactions that are not discernible using other methods.

X-ray crystallography is reliant on the ability to grow crystals, where the constituent molecules arrange in an ordered lattice that is both regular enough and large enough to form a measurable diffraction pattern when interacting with a beam of high energy (short wavelength) photons. Since it is generally energetically unfavourable for large, soluble, macromolecules to pack into an ordered lattice in aqueous solutions, protein crystallisation is a rare event. Finding the correct combination of protein concentration, temperature and added chemical components to make this process favourable takes much trial and error (3). As a consequence, protein crystallographers perform hundreds, if not thousands, of experiments, thereby increasing their chances of success. This, in addition to the difficulty and high cost of producing a pure protein solution, leads to experiments being set up on a small scale (total volume of a few hundred nanolitres), often with the aid of a liquid-dispensing robot.

In a typical vapour diffusion crystallisation experiment, a solution containing the protein is mixed with a ‘cocktail’ of chemical components in order to form a crystallisation drop. Collections of these drops are grouped together on crystal microwell plates that can contain 96 (or multiples thereof) cocktail conditions. Initially, the cocktails are selected from commercial collections that have been collated based upon what has been reported, in the literature, to have been successful for other proteins (4). Once an initial “hit” condition has been found, further optimisation experiments in the chemical space around this set of components can then be used to improve crystal quality and allow scaling-up of the experiments for subsequent studies.

Knowledge of reproducible conditions for protein crystal growth underpins methods such as fragment-based drug discovery (FBDD) (5) and macromolecular room temperature X-ray crystallography (RTX) (6, 7). In FBDD, high numbers of consistent crystals are required for soaking experiments in which large libraries of low molecular weight compounds are trialled as binders to the protein in question. These compounds are added to crystallisation drops after the crystals have grown and, after a set time period, X-ray diffraction data is collected for each combination of compound and crystal. This allows discovery of compound class, binding mode and binding location for compound series that can be subsequently developed into drug leads. For RTX experiments, however, large numbers of smaller crystals are grown in the same droplet or vessel and data collected across multiple crystals is often merged together. Growing these crystals can be achieved using a variety of methods, for example batch crystallisation (8). RTX enables the study of proteins under near-physiological conditions, giving insight into protein dynamics and ligand binding in an environment that is free from cryoprotectants and crystal imperfections caused by the freezing process itself (9).

Regardless of the end goal of the experiment, protein crystallisation trials can be monitored periodically, either manually or via use of a robotic imaging microscope. In the case of vapour diffusion experiments, these robotic “imagers” incubate the experimental plates of crystallisation drops, identified by a unique barcode, and extract them from storage using a motorised arm before capturing images of the drops using a digital microscope camera. The collection of these images (or micrographs) happens on a schedule of predefined timepoints that may cover a period of days or months. The combination of a large number of experiments (drops) and a number of imaging timepoints leads to the creation of hundreds of microscope images that need inspection either by an expert or by an automated system that is able to achieve the accuracy of an expert.

Once crystals have been grown, the subsequent process of data collection for macromolecular X-ray crystallography (MX) has become increasingly automated, particularly at synchrotron experimental stations. In the case of single crystal experiments, crystals are first mounted onto standardised pins (normally by humans) and then robotic arms are used to change samples that are stored in cryo-cooled dewars (10–13), this removes the need to approach the X-ray beam shutter during allocated experimental time and perform subsequent safety checks before opening the shutter again. This, when coupled with faster, more sensitive detectors, leads to higher sample throughput as well as the ability to collect data remotely. To aid this automation, a number of semias well as fully automated solutions have been created to centre the crystal in the X-ray beam, these include X-ray-based centring approaches (14) and image based approaches (15), some of which use deep learning (16, 17). For RTX, a different approach can be taken, with data collection taking place in situ within the environment where the crystals are grown (18–20) this removes the need for manipulating the samples which, in turn, reduces the need for human intervention to collect data.

At DLS, the automated VMXi beamline (18) facility allows RTX experiments to be run in a fashion that minimises the need for any intervention by the scientific users of the beamline in the data collection process. Crystallisation experiments are set up by a liquid-handling robot and the resulting crystallisation microplates are stored in an imaging incubator that is located at the beamline itself. The plates are then imaged periodically on an predefined schedule. Firstly crystal location coordinates for data collection are determined from these images and recorded in the ISPyB laboratory information management system (LIMS) (21). Then, after the user has selected from the list of points and chosen a recipe for data collection, the crystallisation plate is automatically transferred to the beamline imaging hutch. Data collection is performed in situ, with the microplate mounted directly on a goniometer. Any resulting diffraction patterns are then processed by way of automated software pipelines that include the capability to merge data sets from multiple crystals (22). Prior to the work described in this study, both the classification of experimental outcomes and the identification of coordinates for data collection required the scientific user to log into the SynchWeb interface to ISPyB (23) and browse through all of the microscope images before marking the location of points for data collection manually; a time-consuming task.

At the XChem facility, also located at DLS, FBDD campaigns are run on proteins of interest using hundreds of low molecular weight compounds (fragments) (5). Rather than co-crystallising the compound with the protein, many crystals of the same protein are grown in the absence of compound and the fragments are soaked into the crystals afterwards. Generally, after a compound is dispensed into one drop that contains one or more crystals, the molecule is allowed time to diffuse into the crystal lattice before the crystals are manually mounted on pins and cryo-cooled for X-ray data collection. The compounds are dispensed in the form of high concentration solutions in dimethl sulfoxide (DMSO) and precisely targeted into the crystallisation drop using acoustic dispensing. The targeting location is manually determined upon inspection of a micrograph of the crystal drop by the scientist, who will try to choose a point within the drop that is far enough from crystals of interest to prevent damage, either from a high local concentration of DMSO or from the physical stresses associated with bombardment of the crystal with a drop of foreign liquid.

In this study, alongside providing source code and training data, we describe the creation of the two deep learning tools for the DLS VMXi beamline that simplify the task of browsing images and selecting coordinates for data collection. These tools can potentially replace the need for any user intervention at all, thereby removing a major bottleneck in the workflow. The first is an image classification network given the name CHiMP (Crystal Hits in My Plate) Classifier which outperforms the best classifier network from the literature (24) on our in-house images. The second is an image object detection and instance segmentation network, named VMXi CHiMP Detector. Additionally, we describe a third software tool, XChem CHiMP Detector, an image object detection and instance segmentation network that allows automated calculation of coordinates for acoustic dispensing of compound solutions into crystallisation drops. XChem CHiMP Detector facilitates automated dispensing of high concentration fragment compounds for FBBD campaigns that can involve thousands of compound and crystal combinations.

### 1. Background on the automated classification of experimental images

Over the past 40 years, a wide variety of techniques have been investigated to automate monitoring of crystallisation trials with the methodology used reflecting changes in hardware and software capabilities during this period. Comparisons of the effectiveness of these analyses is often difficult, with each institution having it own scoring system for experimental outcomes which then leads to an arbitrary number of categories (e.g. different types of protein precipitation, different sizes and classes of crystals, other phenomena such as phase separation or clear drops) that each image is labelled with. In addition, human expert scorers often disagree on the categories assigned to the same set of images, with a reported rate of *≈* 70% agreement between 16 crystallographers in one study (25) and an agreement rate of between 50% and 70% between three researchers in another (26). This disagreement not only reflects the difficulty of categorising a continuum of outcomes (e.g. *Light Precipitate* versus *Heavy Precipitate*) but also the convention of only assigning one label to an image which may represent multiple outcomes (e.g. an image of a drop containing spherulites and contamination from a clothing fibre could be labelled as either *Spherulites* or *Contaminated*).

#### 1.1. Classification Metrics

A range of metrics are used to report the effectiveness of algorithms that automate the classification of images. Accuracy is widely used but can be a misleading metric for unbalanced datasets such as those from crystallisation experiments; images that record successful crystallisation may be a minority class in the raw data, however conversely, they are often the majority class in images that have been scored. This is due to the fact that scientists will tend to label images that contain crystals and ignore those those that do not. Here we will focus on the precision, the recall and the *F*1 measure of correctly classifying images into the *Crystals* class in order to compare classification performance across the literature as well as to assess the performance of the tools described in this study. This class is a superset of all categories of crystalline outcomes excluding crystalline precipitate.

Precision is defined as:

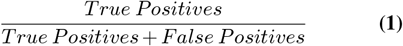

Recall is defined as:

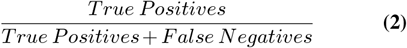

The *F*1 measure is the harmonic mean of precision and recall:

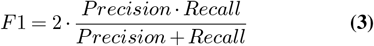

As shown in Equation 1, precision is the ratio of correctly predicted positive observations to the total predicted positives, which means that when a methodology with a high precision score predicts a positive result, it is likely to be correct. A high precision can also be obtained when the method is conservative in its positive predictions, missing the true positives in order to avoid false positives. The recall metric (Equation 2), however, measures the fraction of correctly predicted positive instances retrieved by the method (the sensitivity). Methods with higher recall values capture a larger portion of the true positives, minimizing false negatives. There is a trade-off between optimising a method for each of these two measures, with a more precise method missing some actual positive instances resulting in a lower recall, and a method with a high recall capturing more false positives along with the true positives, reducing precision. The *F*1 score (Equation 3) is a harmonic mean of precision and recall and balances both measures in one metric.

#### 1.2. Previous investigations of automated classification

The first investigations applied image convolutions to enhance vertical and horizontal edges (27) before further interpretation that used a nearest neighbour line tracking algorithm or a Hough transform (28) for detecting straight lines (29). Feature extraction from images coupled with statistical techniques such as linear discriminant analysis (30) (giving a precision 0.24, recall 0.66 and *F*1 0.35 for *Crystals*) or self organising feature maps (31, 32) were later explored.

Several studies focused on extracting image texture information for feature generation (25, 33–35). Machine learning classifiers were trained using these features. For example Cumbaa and Jurisica (34), used 1,492 features to train a 10 category random forest ensemble that achieved a classification precision of 0.64, recall of 0.71 and *F*1 of 0.67 for the *Crystals* class. Another method involved convolution with a bank of filters to assign “texton” (texture prototype) values to pixels, creating histograms of texton frequencies (35). These histograms were used to train a random forest ensemble and the posterior probalility output by the ensemble was used to rank images based on the likelihood of containing “interesting” features, aiding scientists in efficient image selection. The rise of convolutional neural networks (CNNs), especially deeper models (with more trainable layers) has removed the need for extraction of human-designed features since the model is able to adapt or “learn” new convolutional kernels which can extract appropriate features. The pioneering CNN for classifying crystallisation trial images, CrystalNet was a seven layer custom network (36). It was trained on a set of 68,155 images (37) in order to classify images into 10 classes. The resulting precision, recall and *F*1 metrics on the validation set were 0.81, 0.77 and 0.79 respectively for the *Crystals* class. The subset of data used for training consists of images from microbatch-under-oil experiments where three experts independently agreed on the image class. This differs from a real-world dataset where image class is often ambiguous.

In a subsequent investigation by Ghafurian et al. (38), CrystalNet was tested on an internal dataset of 486,000 images at Merck. They found that the accuracy of the model was 73.7% on this data, as opposed to the 90.8% reported in the original study. The authors then went on to train a number of different CNN architectures on their dataset, again using 10 image categories. During training, they compensated for the class imbalance; images containing crystals were a minority class with nine times more images that did not contain crystals (since successful crystallisation is a rare event). This compensation was done using image augmentations to increase the size of their training data, generating images in inverse proportion to the size of each class of image. The final training dataset had over a million images. They found that a deep ResNet model (39) with 56 trainable layers gave the best results with a recall of 0.94 when classifying the test set of *>* 200,000 images.

The next, and arguably the most high profile, CNN to be created for crystallisation micrograph classification was that from the Machine Recognition of Crystallization Outcomes (MARCO) initiative (24). In this work, a customised variant of the 42 layer Inception-v3 CNN architecture was trained on a set of 415,990 images that were collected from a number of organisations including GSK and Merck. Rather than a 10 class classification system, the MARCO network outputs one of 4 labels namely *Crystals, Clear, Precipitate* or *Other*. For the *Crystals* class, MARCO achieves a precision of 0.94, a recall of 0.91 and an F1 measure of 0.93 on the validation set of images. In addition, the authors also created a test set of images of sitting drop experiments solely from their institution that had been hand-scored by an expert. In this set of images, the network achieved a precision of 0.78, a recall of 0.87 and *F*1 score of 0.82. MARCO has been seen as the state-of-the-art in recent years and has been incorporated into commercial software for crystal plate imagers (40).

The MARCO dataset is openly available (41) and has been used to train and evaluate the models in other studies. One approach used two CNNs (42): a U-Net model (43) for segmenting the liquid droplets followed by an Inception-v3 architecture CNN for classifying experimental outcomes. The classifier network was pre-trained on a subset of the ImageNet database of images (44) which consists of 1.2 million images from 1000 mutually exclusive classes. Pre-training in this fashion results in a model that has already learnt a feature representation for extracting information from images. The weights from this network are then fine-tuned to perform a new classification task; the advantage of this strategy over training a model with randomised weights is fewer training epochs are required and there is less chance of over-fitting the model when the training dataset is small. The dataset used in this particular study was a small subset of the MARCO dataset (150 to 450 images for the U-Net and 99 images for the Inception-v3 network). The authors concluded that using their two CNN approach was superior to using one classifier network, however, the final precision and recall metrics for classifying the images are not given.

In another study, an EfficientNet CNN architecture (45) was trained on the full MARCO dataset (46). In this case, the same validation set was chosen as used by (24) but a separate test set of images was also created, comprising of 10% of the original training set. The images were classified into the same four categories as MARCO. On the validation set they reported a precision of 0.87, a recall of 0.87 and an *F*1 score of 0.87 for the *Crystals* class. On their test set they reported a precision of 0.95, a recall of 0.97 and an *F*1 metric of 0.96 for *Crystals*. This finding is somewhat unexpected, given that this data originally came from the same pool of images as the training and validation sets. For comparison, the authors then used the original MARCO model to classify the same test set (which had formed part of the training set for this particular model) resulting in a precision of 0.94, a recall of 0.91 and an *F*1 of 0.92 for *Crystals*; these metrics correspond with the published results from (24). The rationale behind utilising this test set for comparisons rather than the original validation set is unclear.

As part of a study which carried out a further investigation into the structure of the MARCO dataset, Rosa et al. (47) trained a ResNet50 network on the data. They achieved a slight improvement in overall accuracy (94.63% vs the original MARCO score of 94.5%). The authors also found that supplementing the MARCO training data with local images (images from their own equipment in their institution) helped improve the accuracy by as much as 6% when classifying these local images as opposed to using the MARCO dataset alone. In addition, they examined the reasons behind the poorer performance of models trained on the MARCO data alone when classifying images from sources that are not part of the original data. These reasons include data redundancy (duplicate images), the resolution of images included in the data, mislabelling of images, plate and imager types and the inclusion of a small number of images from Lipid Cubic Phase (LCP) experiments; these particular images can have a dramatically different appearance to images of standard sitting drop vapour diffusion experiments.

In their study, Milne et al. (26) fine-tuned four different architectures of CNN on their own dataset of 16,317 images. The classifiers were pre-trained on ImageNet and were chosen to be trainable with more limited computational resources. The authors found that they achieved better performance by only using their own images and not supplementing them with those from the MARCO dataset. Each model was found to have its own strengths and weaknesses at categorising images into the 8 categories that they had defined. For example, for images containing crystals, the Xception architecture (48) was found to have the highest precision, at 0.96, however the recall was 0.75, giving an *F*1 of 0.84. As a result of this, they opted to combine multiple classifiers into an ensemble of models, where majority voting was used, to achieve an accuracy comparable to MARCO. This was possible even when excluding the Xception network and just using the other three architectures, namely Inception-v3, ResNet50 and DenseNet121 (49).

Thielmann et al. (50), trained four different CNN architectures on a dataset generated by their own robotic imaging microscopes. These were AlexNet (51), VGGNet (52), ResNet50 and SqueezeNet (53). They chose to adopt an approach similar to (42) and first trained a U-Net network to detect the experimental droplets. After detection, the droplet region of interest (ROI) was further subdivided into image patches that were used for training a separate CNN model to classify the images into 12 classes. Their initial dataset size was 33,872 images and their process of subdividing the images led to *>* 324,000 patches (the authors did not state the split between training and validation sets). In order to classify the images it was again necessary to divide the image into patches and then classify each patch individually before arriving on a class for the parent image using a ranking system. They achieved the best overall success with the SqueezeNet architecture although AlexNet gave the highest success for the *Crystals* class.

### 2. Background on crystalline object detection in images

Despite many years of research effort dedicated to classifying experimental micrographs into categories, it is only more recently that the possibility of detecting the locations of the protein crystals and/or the drops within the images has been investigated. Object detection techniques define a bounding box for the entity in the image and assign a class label, giving an approximate measure of size as well as the (*x, y*) coordinates. For more accurate sizing of entities in the image, instance segmentation can be used. Instance segmentation is the process in which individual objects in an image are separately identified before all the pixels in the object are given a label. This allows the counting of, as well as measurement of, crystals from images to give size distributions.

Crystal object detection and instance segmentation have been investigated most thoroughly in the context of batch experiments on proteins and small molecules in the field of Chemical Engineering. The use of reproducible procedures means that the resulting crystal morphology is often known which allows model-based approaches to be used (54, 55). The size distributions measured by these techniques can be used to characterise the end products of industrial processes as well as for monitoring the crystallisation process over time.

#### 2.1. Object detection metrics

To assess the performance of object detection networks, the metric of mean Average Precision (mAP) is most commonly used. The basis of the mAP metric is the precision and recall (as described in Section 1.1, Equations 1 and 2) as used for classification tasks. However, in the context of object detection where a bounding box is predicted for each object, the definition of what constitutes a correct classification is dependent on the degree of overlap between the predicted bounding box and the actual “ground truth” bounding box. This degree of overlap is measured using the Intersection over Union (IoU) metric which is defined in Equation 4. When considering instance segmentation, IoU is calculated in the same manner but with intersection and union of the segmented areas as the equation terms.

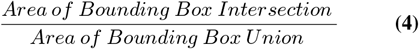

In order to calculate mAP, it is first necessary to plot a precision/recall curve for each class. This is a plot of how these two metrics change at different confidence thresholds for a classifier. The average precision, defined as the area under this curve, is then calculated for a range of IoU thresholds (although in some cases at just one IoU threshold). The mean Average Precision is the mean of these calculated average precision values when also averaged across all the classes detected. In addition to defining the average precision by IoU threshold, evaluation can be split by object size in order to allow comparison of performance on “small” (*<* 32^2^ pixels) versus “large” (*>* 96^2^ pixels) objects, for example.

Due to the complexities of mAP, to compare scores with one another it is important to be sure that the mAP has been calculated across the same range of IoU thresholds and also calculated on the same model output (bounding boxes versus segmentation masks). Although a higher mAP score means more accurate detection and localisation of objects in images, there is no notion of what constitutes a good score. In all but one of the previous studies described here, no metrics were reported. In our study we report the mAP achieved so that it may serve as a benchmark for future work.

#### 2.2. Previous investigations of automated crystal detection

Gao et al. (56) trained a network to perform object instance segmentation of crystals of L-Glutamic acid. The Mask-R-CNN architecture that they used (57), has a backbone network (often ResNet50 or similar) that extracts features as part of a feature pyramid network (FPN). The levels within the FPN contain features at different resolutions allowing the network to be able to detect objects at different scales in the image. The feature maps from the FPN are then passed to a region proposal network (RPN) which predicts bounding box proposals for regions of interest within the image. The underlying features are aligned with the region proposals before being passed on to parallel network components which output bounding box coordinates, a class label for the object and a segmentation mask. The trained network was able to predict instance segmentation masks for both *α* and *β* crystal forms and allowed crystal size distributions for both populations to be determined from images. No performance metrics are given for this network, however.

Bischoff et al. (58) trained a Single-shot Alignment Object Orientated Detection network (S^2^A OOD-Net) (59) on simulated images of protein crystals in suspension. Rather than outputting bounding boxes of a fixed orientation, which was previously standard for object detection networks, this network is able to orientate the boxes to achieve the best fit to crystals. The synthetic images were generated using ray-tracing algorithms with the size and alignment of the crystals determined from a statistical distribution and the crystal shape chosen at random from a collection of 1851 models. This resulting training set contained 332,558 images. The model was evaluated by calculating size distributions of crystals in experimental images of *Lactobacillus kefir* alcohol dehydrogenase which had been grown in stirred tank batch reactors. No metrics were calculated for precision of detection (such as mean adjusted precision).

In order to apply object instance segmentation to sitting drop protein crystallisation micrographs, Qin et al. (60) used a Mask-R-CNN with a ResNet101 backbone network. This backbone had been pre-trained on the Common Objects in Context (COCO) dataset (61). The COCO dataset contains 2.5 million labelled segmentation instances and utilising a model pre-trained on this data has the same advantages as utilising classification networks pre-trained on the ImageNet dataset. The images used for training their network were taken from the MARCO dataset but since this data is annotated solely with classification labels, instance segmentation masks had to be created first. The masks were created manually on the images using a drawing tool called Labelme (62). The authors do not disclose how many images were annotated or the size of their training set but they did report a mAP score of 0.47 (calculated on 100 images at an IoU threshold of 0.65) and 0.70 (calculated on 10 images at an IoU threshold of 0.5). An improvement in the performance of their model was achieved when including an image pre-processing step, namely the Contrast Limited Adaptive Histogram Equalization (CLAHE) algorithm (63). A network trained on images which were not processed using CLAHE had a lower mAP score of 0.30 at an IoU threshold of 0.65 and 0.67 at an IoU threshold of 0.5. This localised image enhancement potentially allows crystals in the shadow areas at the edges of drops to be discerned more easily and also could reduce reflection artefacts from the surface of the drops.

##### Justification for creating new, in-house, tools

To enable further automation of the VMXi robotic in situ macro-molecular crystallography beamline at DLS, automatic categorisation of experimental micrographs alongside detection of crystal positions for fully-automated data collection was desired. The initial solution implemented for categorising the images was to use the MARCO Classifier network (24), as described in section 1.2. The network was deployed in November 2019 and tasked with classifying all images collected by the Formulatrix Rock Imager machines at the experimental facility. The predicted label probabilities and resulting image labels were added into the ISPyB LIMS database and then could be viewed on a schematic plate overview in the SynchWeb interface by beamline users (shown in Figure 8, Supplementary Section 1). After an initial period of use, however, the feedback from those using the new feature was mixed, with mistrust that the images were being classified correctly. As a result of this, an experiment was then performed in order to evaluate the performance of the MARCO Classifier on micrographs captured in-house. This work found that the *F*1 score for the MARCO network when categorising images from VMXi as *Crystals* (0.76) was slightly lower than reported for the same network on a independent test set of images (0.82), (24). The main finding, however, was that the MARCO Classifier tended to classify images as *Precipitate* at the expense of other classes, with a recall of 1.0, a precision of 0.48 and a false positive rate of 37%. This bias toward categorising images as *Precipitate* is understandable since *≈* 48% of the MARCO training dataset is made up of images that were given this label and these images outnumber the images labelled as *Crystals* by a ratio of *≈* 4:1. The methodology and more detailed results of this experiment are described in Supplementary Section 1 subsection 1. As was later confirmed by Rosa et al. (47), we hypothesised that training with local images would be needed in order to achieve better classification performance since the images in the MARCO training dataset come from different imagers and the experiments are set up in different plates to those used in our facility. Difficulties associated with adding training to the existing MARCO model, along with advances in the field and the need to perform object detection as well as image classification, led us to carry out experiments on training our own networks.

Automation for finding the (*x, y*) coordinates of any crystals present in the experimental images was desired for two different use cases. In the first, on the VMXi beamline, human intervention was required to locate each crystal and add it to a queue for data collection. At the time, this was done using a web-based point and click interface in SynchWeb which then added the coordinates to the ISPyB LIMS system. In the second case, as part of the XChem project at DLS, high throughput fragment-based drug discovery workflows required acoustic dispensing of low molecular weight compounds into the experimental drops after crystals had grown. High levels of the solvent DMSO in the direct vicinity of the crystals can disrupt their structure and ability to diffract. Because of this, dispensing of the compounds is targeted within the drop but in a region away from the crystals to be mounted for data collection, thereby allowing the compound and associated DMSO to diffuse through the drop gradually towards the crystals of interest. At the time, this targeting step was done via a manual point and click interface in a modified version of TeXRank software (35) but automation of this targeting was desired in order to increase throughput of screening pipeline.

## Methods

The procedures used to collate the data and to train and evaluate the classification as well as the detection networks are described here.

### 3. Creating the CHiMP Classifier models

The data used for training crystal micrograph classification networks comprised two sets of images, the first, henceforth referred to as the VMXi Classification Dataset is described in more detail in Section 3.1. The second set of images was obtained from the Machine Recognition of Crystallisation Outcomes (24) project and is henceforth referred to as the MARCO Dataset (this dataset has been introduced in Section 1.2 and is further described in Section 3.2).

The initial work on classification networks used the VMXi Classification Dataset alone. This resulted in a ResNet50 classification model given the name of CHiMP (Crystal Hits in My Plate) Classifier-v1 (more detail about the training of this classifier is given in Section 3.3). This network was put into production on the VMXi beamline for a period of three years. Although the feedback from use of this classifier was positive (to the extent that image classification by the MARCO classifier was eventually switched off), further work was started to investigate replacing the network. This was done for two main reasons: Firstly, there was a software dependency on version 1 of the *fastai* package. The second version of the package had an API that was not backwards compatible and development on the first version was stopped. Secondly, although the ResNet50 model weights could have been transferred to a purely PyTorch ResNet50 implementation, significant progress had been made in CNN architectures in the meantime. A more modern model architecture, namely ConvNeXt (64) was chosen. As well as utilising increased kernel sizes, ConvNeXt networks use some innovations commonly found in Vision Transformer (ViT) models, such as reduced use of activation functions and normalisation layers, and applies these modifications to a CNN in order to improve performance. The ConvNeXt *Tiny* version of the model was chosen because it has a similar number of parameters to ResNet50. Once trained, this new model, named CHiMP Classifier-v2, was put into production on the VMXi beamline in March 2023 (more detail about the training of this classifier is given in Section 3.4).

#### 3.1. Curation of the VMXi Classification Dataset

Details of the collection and manual categorisation of the images at the VMXi beamline experimental facility are given in Supplementary Section 2, Subsection 1.

At VMXi, a 10 class classification system is used for labelling the outcomes (see Table 1). The initial set of images that makes up the VMXi dataset were found by querying the ISPyB LIMS for images which had associated scores and that had been collected up to the query date (March 2020). The set of images was also further restricted to those from experiments belonging to beamline users who were not from commercial/industrial organisations in order abide by intellectual property restrictions and agreements. This resulted in a set of 18,782 images. In order to remove redundancy in the data caused by inclusion of similar images from multiple inspections of the same subwell, the data were grouped by subwell and the inspection numbers analysed. The majority of the subwells only had one image with an associated score (11,167 images), these were included in the final dataset. In the cases where subwells had images with scores for more than one inspection, the image and associated label from the second of these inspections was added to the dataset (2,784 images). This resulted in a final dataset size of 13,951 images.

**Table 1.** Mapping of VMXi image scoring labels to a four-label system.

| Score | VMXi Label | Mapped Label |
| --- | --- | --- |
| 0 | Clear | Clear |
| 1 | Contaminated | Other |
| 2 | Light Precipitate | Precipitate |
| 3 | Heavy Precipitate | Precipitate |
| 4 | Phase Separation | Other |
| 5 | Spherulites | Other |
| 6 | Microcrystals | Crystals |
| 7 | 1D Crystals | Crystals |
| 8 | 2D Crystals | Crystals |
| 9 | 3D Crystals | Crystals |

The labels assigned to the images were mapped from the 10 in-house categories to a four class system (as used by MARCO): *Crystals, Precipitate, Clear, Other*. This mapping can be seen in Table 1. The breakdown of the number of images in each category at this stage can be found in Supplementary Section 2 Table 8.

This dataset was used as the starting point for fine-tuning a ResNet50 network that had been initialised with weights from training on ImageNet. More details of the training method can be found in Section 3.3. During the initial rounds of training it became apparent that the training labels associated with the images were not always accurate, so a strategy to clean the image labels was implemented. Details of this method can be found in Supplementary Section 2, Subsection 2. In total 904 labels were changed during this process (6.5% of those in the dataset), an overview of the changes broken down by image class can be seen in Supplementary Section 2 Table 8. After cleaning, the data was again split at random into a training set comprising 80% (11,161) of the images and a validation set of 20% (2,790 images). The final number of images in each category can be seen in Table 2. Example images from each class can be seen in Figure 1. The dataset is available to download from doi:10.5281/zenodo.11097395.

**Table 2.** Breakdown of image classes in the cleaned VMXi Classification Dataset.

| Label | Training Set | Validation Set | Total |
| --- | --- | --- | --- |
| **Crystals** | 6,752 | 1,654 | **8,406** |
| **Clear** | 1,247 | 282 | **1,529** |
| **Precipitate** | 2,430 | 671 | **3,101** |
| **Other** | 732 | 183 | **915** |
| **Total** | **11,161** | **2,790** | **13,951** |

**Fig. 1.**
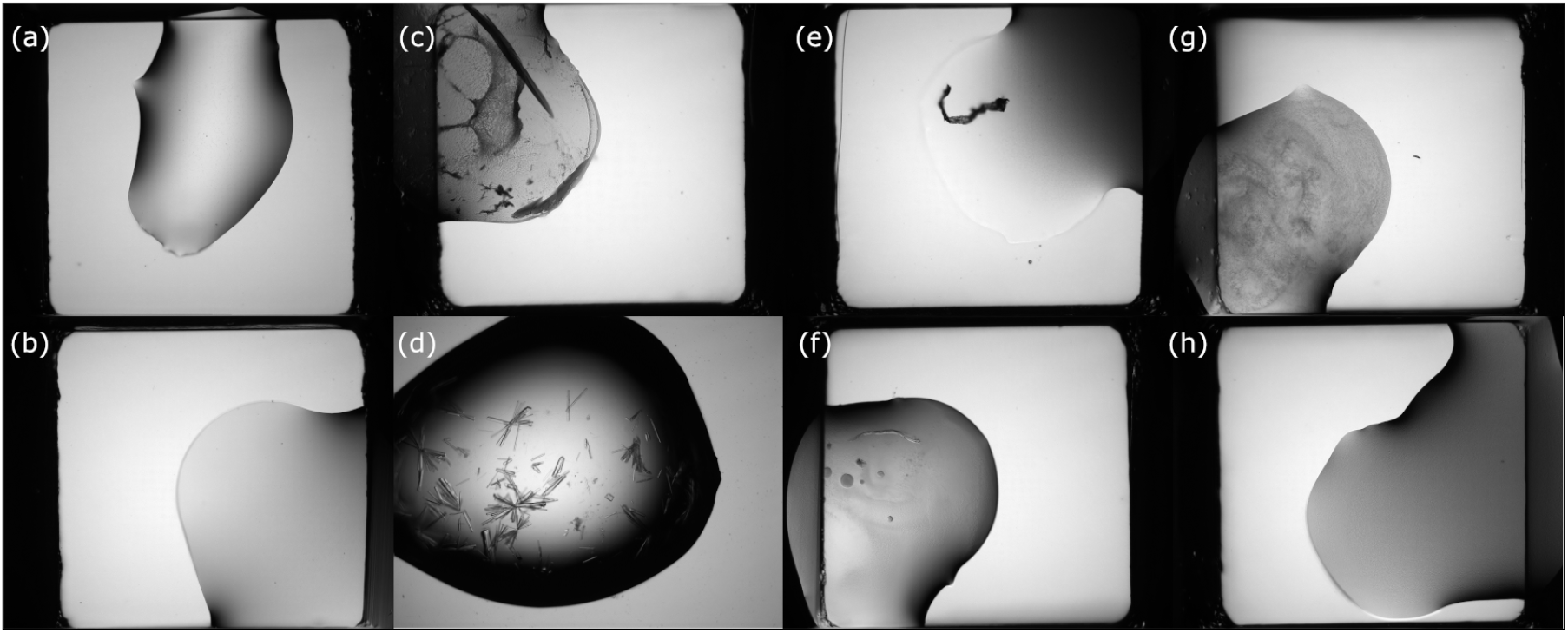
Examples from the VMXi Classification Dataset. Two images from each category selected at random from the training dataset. (a, b) Images from the *Clear* class.(c, d) Images from the *Crystals* class. (e, f) Images from the *Other* class. (g, h) Images from the *Precipitate* class.

#### 3.2. The MARCO Dataset

As described in Section 1.2, the Machine Recognition of Crystallization Outcomes (MARCO) initiative (24) collated 493,214 labelled images from five different organisations to create a dataset for training their CNN classification model. Ambiguous labels were cleaned in a similar fashion to that described for VMXi classification datset in Supplementary Section 2, Subsection 2, with the top 5% of images (ranked by classification loss) revisited by expert crystallographers. In this 5% of the images, 42.6% were relabelled, which highlights the “label noise” present in the dataset.

After cleaning, the authors divided the data into training and validation sets comprising *≈* 90% and *≈* 10% of the images respectively. In their paper, Bruno et al. (24), give a table summarising the breakdown of image classes in the MARCO dataset (with 442,930 training and 50,284 validation images) however, the dataset that they used for training is described as having 415,990 training images and 47,062 validation images. The numbers of images in the dataset that they released publicly differs slightly again. A summary of the number of images in each category in this publicly released set is given in Table 3 since this is the one used in our investigations. A detailed exploration of the MARCO dataset can be found in (47). Example images from each class can be seen in Figure 2.

**Table 3.**
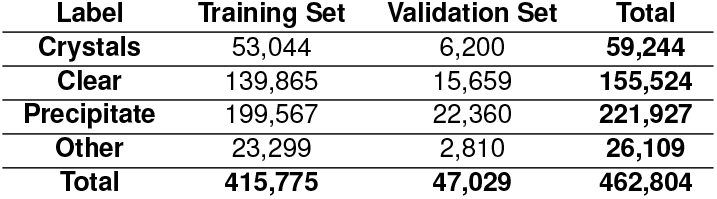
Breakdown of image classes in the publicly released MARCO Classification Dataset. Available from https://marco.ccr.buffalo.edu/download

| Label | Training Set | Validation Set | Total |
| --- | --- | --- | --- |
| **Crystals** | 53,044 | 6,200 | **59,244** |
| **Clear** | 139,865 | 15,659 | **155,524** |
| **Precipitate** | 199,567 | 22,360 | **221,927** |
| **Other** | 23,299 | 2,810 | **26,109** |
| **Total** | **415,775** | **47,029** | **462,804** |

**Fig. 2.**
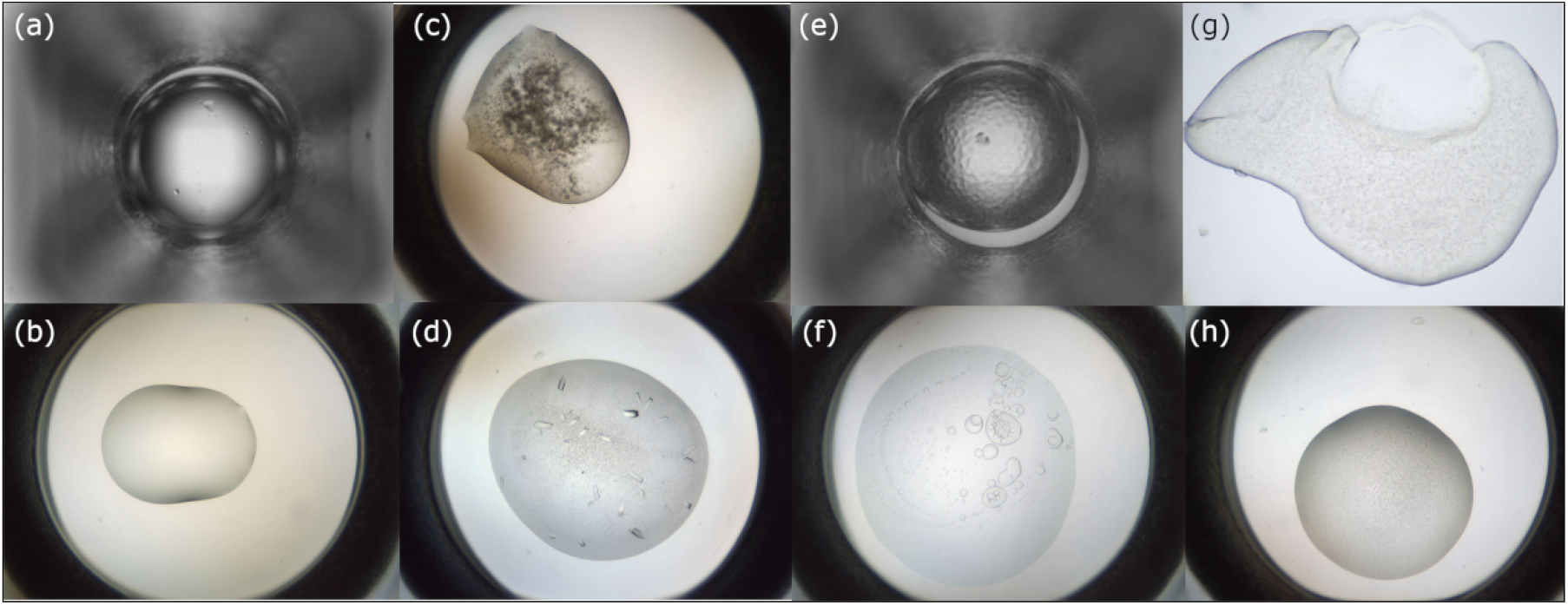
Examples from the MARCO Dataset (24). Two images from each category selected at random from the training dataset. (a, b) Images from the *Clear* class. (c, d) Images from the *Crystals* class. (e, f) Images from the *Other* class. (g, h) Images from the *Precipitate* class.

#### 3.3. Training of CHiMP Classifier-V1

This initial work on classification networks used the *fastai* Python library (65), which is a high level interface to the PyTorch machine learning framework (66).

A ResNet50 convolutional neural network (CNN) (39) that had been initialised with weights from a model trained on the ImageNet database (44) was the starting point for our experiments. This model was fine-tuned on the VMXi Classification Dataset (described in Section 3.1) in a number of phases with increasing image dimensions. To prevent overfitting to the training data (to make up for the relatively small number of images available), during each phase of training, the training set of 11,161 images was augmented using a randomised range of flips, rotations, warping and zooming as well as brightness and contrast adjustments. Before passing the images through the network, they were reshaped (or “squished”) to a square, rather than cropped, this was done to avoid objects at the edge of the image, such as crystals being excluded. In addition the image histograms were normalised to match those used by the ImageNet database. In order to make up for the label imbalance in the VMXi classification dataset, (where the images labelled as *Crystals* make up *≈* 60% whereas those in the *Other* class make up just 6.6% of images), an oversampling method was used. This method sampled the image classes in the dataset in inverse proportion to their frequency, thereby counteracting this imbalance. In total the model was trained for 31 epochs with a batch size of 64, the final model had a cross entropy loss of 0.115 for the training set and 0.364 for the validation set. Training was carried out on an NVIDIA Tesla P100 GPU with 16GB of VRAM and took around 6.5 hours of computation time. Further details of the training strategy can be found in Supplementary Section 2, Subsection 3. Metrics on the validation set of images from the VMXi Classification Dataset are given in Section 5.1. Evaluation of the performance of this model on an independent test set of images from the VMXi facility is given in Section 5.2.

The network was deployed alongside MARCO to classify all experimental images that were produced by the imagers on the VMXi beamline. The classification labels were inserted into the ISPyB LIMS and the results were displayed on a plate overview schematic in the SynchWeb interface, (shown in Figure 3).

**Fig. 3.**
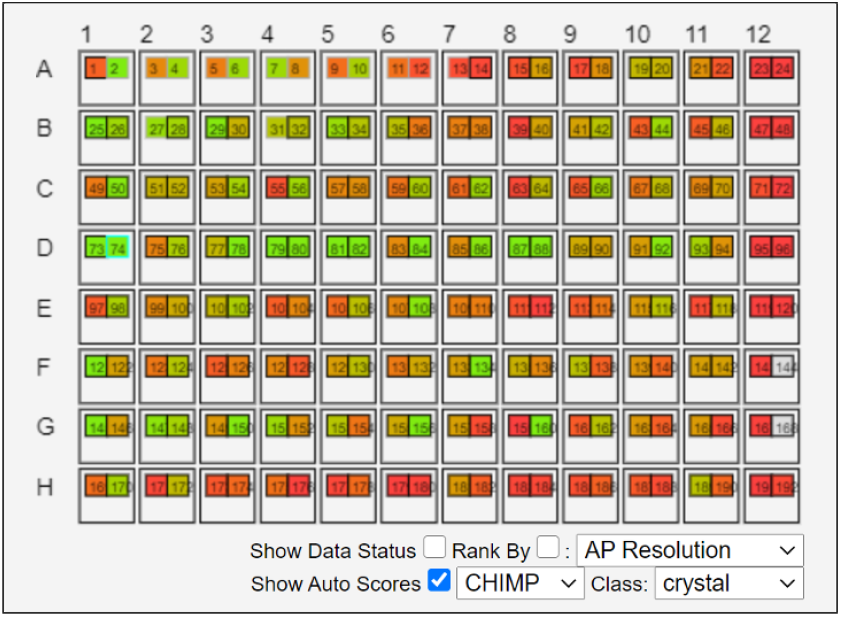
Schematic overview of classification outputs from the CHiMP Classifier-v1 CNN, as displayed in the SynchWeb interface (23) for users of the VMXi beamline at Diamond Light Source. The 96 plate wells are represented as larger boxes with the two subwells containing crystallisation droplets shown as smaller boxes within. Each subwell is assigned a colour on a gradient from red to green that represents the probability of the associated image being assigned to the class *Crystals* with green representing a high probability.

#### 3.4. Training of CHiMP Classifier-V2

In a similar manner to the work on the first version of the classifier, the starting point for training was a model initialised with weights from ImageNet pre-training. This model was then fine-tuned using custom PyTorch training routines. Unlike version 1 of the CHiMP Classifier, in this case, the first stage of training was performed using the MARCO dataset (described in Sections 1.2 and 3.2), and then the final training stages used the VMXi Classification Dataset (described in Section 3.1).

The pre-trained ConvNeXt-Tiny network was obtained from the *PyTorch Image Models* library (67). The standard one-layer classification head of the CNN was replaced with a two layer fully-connected network with batch normalisation layers and dropout of 0.4 on the last layer, rather than using a standard, single fully connected layer. An initial training epoch was carried out with all convolutional parameters frozen; training only this classification head to start with.

A number of training phases were implemented, with increasing image dimensions in each phase before final training epochs on images with dimensions of 512 × 512 pixels. Images were resized to the required input dimensions by re-scaling followed by padding in order to ensure that objects such as crystals at the edge of the image were not removed by cropping. Image augmentations that were used on the training set included random flips and rotations, optical distortions, brightness and contrast alterations, blurring and contrast-limited adaptive histogram equalisation (CLAHE). The imbalance in the number of images in each category in the dataset was counteracted by sampling the data in inverse proportion to the frequency of each category. After each training epoch, the model was saved and used as the model for the next epoch only if the overall classification accuracy on the validation set had improved in comparison to the previous best model. In order to prevent over-fitting, an early stopping safeguard was used, which terminated model training if the validation accuracy stopped improving after a predefined number of epochs. As for version 1 of the classifier, the optimiser for model parameters was AdamW (68) and cross entropy loss was used. The learning rate was again cycled during training on a “1cycle” schedule (69).

Further details of the training phases can be found in Supplementary Section 2, Subsection 4. After 12 epochs of fine-tuning on the MARCO Dataset, the cross entropy loss for the training set was 0.201 and for the validation set was 0.214. After a further 24 epochs of fine-tuning on the VMXi Classification Dataset, the final cross entropy training loss was 0.421 and the final validation loss was 0.522. Metrics on the validation set of images from the VMXi Classification Dataset are given in Section 5.1. Evaluation of the performance of this model on an independent test set of images from the VMXi facility is given in Section 5.2.

This new model, named CHiMP Classifier-v2, was put into production on the VMXi beamline in March 2023.

#### 3.5. Creation of test sets of images to evaluate model performance

A test set of images from the robotic imaging microscopes on the VMXi beamline was created and scored by a panel of experts in order to evaluate the classification performance of the networks. At the time of creation of this test set (August 2022), the CHiMP Classifier-v1 network had been categorising all images collected by the imagers for more than two years, with the results inserted into the ISPyB LIMS. The LIMS was queried for scored image labels and returned data for around 780,000 classified images. These classifications were first grouped by inspection number and plate subwell before limiting the set to those with two inspections or more in order to exclude images taken of unusual experiments that were not being monitored over time. The remaining set consisted of 751,540 images from 69,542 subwells.

The groups of images for each subwell represent time-courses over which the experiments were monitored. To prevent multiple similar images from the same subwell appearing in the dataset, one image was selected at random from each of these time-courses in the range of inspection 2 to inspection 7. This population of 69,542 images with associated classification labels was then sampled to create a final dataset of 1000 images with 250 members from each class from the four category system used by CHiMP Classifier-v1; (*Crystals, Precipitate, Clear, Other*). This balancing was done to ensure that the dataset included sufficient numbers of images from each class to be able to draw robust conclusions.

A panel of three experts viewed all images independently and assigned one of the four class labels to each image. Experts did not always agree with each other (described further in Section 5.2). To accommodate this ambiguity in labelling, two subsets of the dataset were created to assess the models against. The first, named the *unambiguous* test set, consisted of the 632 images where all three experts agreed on a label. The second, named the *mostly unambiguous* test set consisted of 949 images where at least two experts agreed on the label. The number of images assigned to each category for each of these datasets can be found in Table 4.

**Table 4.** The number of images in each category for the VMXi Classification Test Sets. Sets were collated after scoring of 1000 images by three experts. **Unambiguous** - images where three experts agreed on a label. **Mostly Unambiguous** - images where two or more experts agreed on a label.

| Label | Test Set |  |
| --- | --- | --- |
|  | Unambiguous | Mostly Unambiguous |
| Crystals | 145 | 179 |
| Clear | 95 | 182 |
| Precipitate | 230 | 356 |
| Other | 162 | 232 |
| Total | 632 | 949 |

### 4. Training Mask R-CNN networks to detect objects in experimental micrographs

Initial experiments used Gradient-weighted Class Activation Mapping (Grad-CAM) (70) to locate regions of the image where CHiMP Classifier V1 was being activated when predicting the *Crystals* class. However, the low resolution of the output maps (16 × 16 pixels) meant that this technique was not able to provide a positional accuracy high enough for our purposes. An alternative approach of training a CNN such as a U-Net to perform segmentation of the crystals from the rest of the image was also considered. The disadvantage of this approach is that a *semantic segmentation* technique such as this would divide the image into two classes, crystals and background but not separate out individual *instances*. Detecting individual crystal instances would be preferable, allowing those that are overlapping or growing in conjoined inflorescences to be detected and targeted individually. For these reasons, use of an object detection CNN was investigated, in this case a Mask R-CNN architecture (described in Section 2.2), which has the advantage of also providing instance segmentation alongside bounding box suggestions, providing the potential to allow more accurate measurement of crystal dimensions and calculation of the centre of mass for targeting.

#### 4.1. Image selection for detection networks

To obtain images to train the detection network for the VMXi beamline, first a random selection of 1000 experimental images from different plate subwells was taken from the set of all images collected up to that point (March 2021), excluding those linked to industrial experiments. These images were then classified automatically using the CHiMP Classification-v1 network before a dataset of 250 images was sampled from this initial selection. The sampling was done with 50% (125 images) coming from those images labelled as *Crystals, ≈* 25% (62 images) labelled as *Precipitate, ≈* 20% (51 images) labelled as *Clear* and *≈* 5% (12 images) labelled as *Other*. Since each image would need to be annotated manually, the size of the training data that could be created would be limited, therefore this split was chosen to prioritise images with crystals present whilst also including those with a range of different outcomes for the network to learn from. These images were then inspected manually and similar images discarded resulting in a final dataset size of 237 images. These images were then scaled down by a factor of two to 1688 × 1352 pixels before having their histograms adjusted using the CLAHE algorithm (63) using the *OpenCV* library (71) with grid size of 12. This was done to enhance visibility of crystals in shadow areas and reduce glare from reflections on droplets.

To obtain images to train the detection network for the XChem Fragment-Based Drug Discovery facility, the integrated database of their RockImager 1000 robotic imaging system (Formulatrix, USA) was queried (October 2022). The images were then grouped by plate subwell and a selection of 1000 images chosen at random from this grouped set. These images were then viewed individually and reduced to set of 350 images that showed a diversity of experimental outcomes, plate types and crystal forms. Theses images, with dimensions 1024 × 1224 pixels, then had their histograms adjusted using the CLAHE algorithm in the same manner as done for the VMXi image dataset.

#### 4.2. Creating image annotations for detection networks

When training a Mask R-CNN model, three sets of label data are needed for every object contained within the image, alongside the image files themselves. These are: (i) Object masks (the outline of each object), (ii) object class labels and (iii) object bounding box coordinates (coordinates of the vertices of a rectangular selection that encloses each object). Manually generating this information for each image can be arduous depending on the image content e.g. in the case where there are hundreds of protein microcrystals in a droplet. In order to generate enough high quality image annotations to train a network, the task of annotating was shared amongst experts at DLS using a custom project on the Zooni-verse web platform (www.zooniverse.org), (72). A Zooniverse workflow was created with drawing tools that could be used to create polygonal masks on experimental droplets and on crystals observed in the images. The set of images to be annotated was uploaded to the platform and each expert annotator given access to the project. Figure 4 shows an example of using this workflow. The platform randomly selects images from the pool of unannotated images that remain and displays them to the expert alongside the web-based annotation tools. Once completed, the annotations can be downloaded from the platform in the form of a Comma Separated Values (CSV) file with the data contained in JavaScript Object Notation format (JSON) strings within this file. Methods were written in Python to extract the data from these strings and convert it into labels, masks and bounding boxes suitable for training a Mask R-CNN network (*available at ?*).

**Fig. 4.**
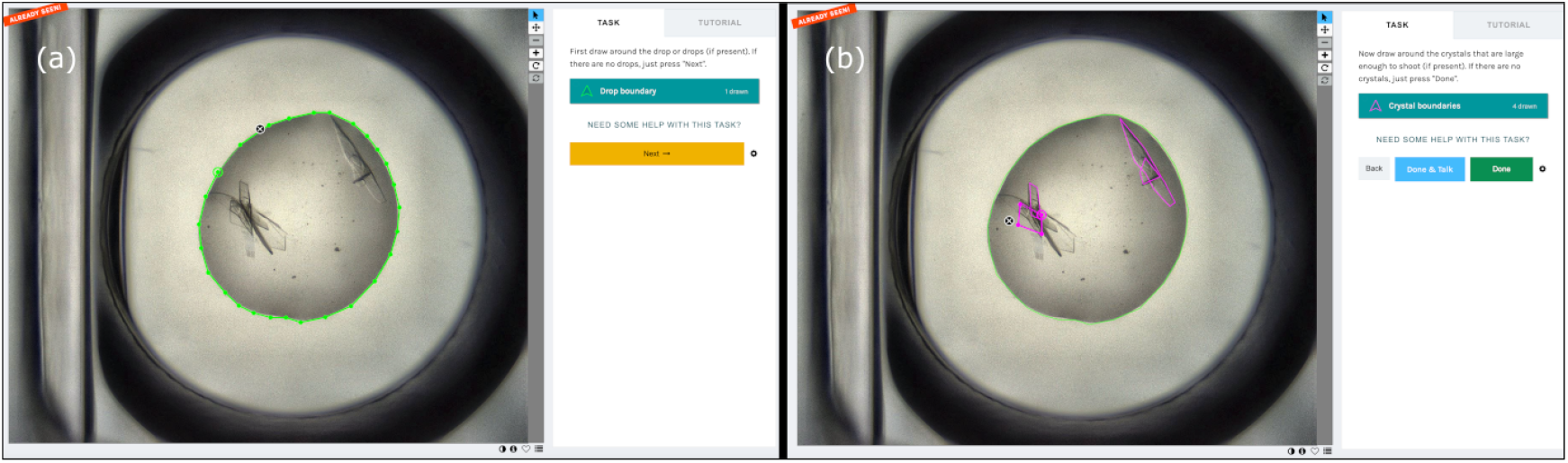
The Zooniverse annotation interface for generating training data for networks that perform object detection and instance segmentation. (a) An example of annotating a crystallisation drop. (b) An example of annotating crystals.

#### 4.3. Training the VMXi CHiMP Detector network

The 237 images and associated annotations comprising the VMXi Detector Dataset were split into training and validation sets with 190 images (*≈* 80%) in the training set and 47 images in the validation set. Since only the positions of the crystals were required, only the labels for the crystals were used in training, this gave better metrics than initial experiments than when including labels for drops at the same time (results not shown). Because of the small size of the training dataset, images (along with associated object masks and bounding boxes) were augmented using random flips and rotations as well as random blurring, sharpening and contrast and brightness adjustments. The model was initialised with weights from a model pre-trained on the COCO dataset. Training was done for 20 epochs. The learning rate was cycled during training using a “1cycle” policy (69) with an average learning rate of 1 × 10^*−*4^. The optimiser used was AdamW. The different components of the Mask R-CNN model use different loss functions (e.g. mask loss, bounding-box loss, classification loss) which are combined into one value named multitask loss for monitoring during training. Training was done with a batch size of 4 on an NVIDIA P100 GPU with 16 GB of VRAM and took 1 hour and 10 minutes to complete. The final multi-task training loss was 0.7749 and the final multitask validation loss was 0.8335. This model was named the VMXi CHiMP Detector network and was deployed onto the VMXi beamline in December 2021. Crystal targeting coordinates are calculated by taking the centroid (centre of mass) of the segmentation masks for each crystal detected. These coordinates are then inserted into the ISPyB LIMS database. The SynchWeb interface for the VMXi facility was altered to display these points and allow them to be queued for data collection.

#### 4.4. Training the XChem CHiMP Detector network

The training of a Mask R-CNN for use on images from the XChem FBDD experiments took place approximately one year after the training of the VMXi CHiMP Detector network. Because of this, the strategy for training was different. In this case, the starting model was a Mask R-CNN that had already been fine-tuned for 20 epochs on both the crystal and drop annotations from the 237 images in the VMXi dataset. This model was then trained further on the 350 images and associated annotations that make up the XChem Detector Dataset. The data was split randomly with 280 images (80%) in the training set and 70 images in the validation set. This training was done in the same fashion as for the VMXi CHiMP detector with image augmentations including random flips and rotations as well as random blurring, sharpening and contrast and brightness adjustments. Once again, a “1cycle” policy was used along with an AdamW optimiser. Training was done for 50 epochs with an average learning rate of 5.2 × 10^*−*4^. The batch size used was 4 and the training was performed on an NVIDIA V100S GPU with 64 GB of VRAM and took 1.25 hours to complete. The final multi-task training loss was 0.7069 and the final multi-task validation loss was 0.6962. This model was named the XChem CHiMP Detector network.

#### 4.5. Using the XChem CHiMP Detector network to calculate compound dispensing positions

Using outputs from the Mask R-CNN trained on images from the XChem project, an algorithm was developed to determine a candidate position in the experimental drops for acoustically dispensing compound fragments. The network creates segmentation masks for drops detected in addition to a mask for every crystal detected. The ideal position for dispensing compound (i) is away from the edge of the drop, yet also (ii) away from where the crystals are located. A position that meets these two criteria was calculated as follows:

The segmentation mask for every crystal detected was subtracted from the drop mask resulting in a single mask which encodes the shape of the drop minus the shape of the union of all crystals in that drop. This binary mask was then processed using an exact Euclidean distance transform. This transform replaces each pixel in the mask with a value denoting the Euclidean distance to the nearest pixel outside the mask, (for more detail on this transform see (73)). After the transform, the pixel with the highest value is that which is furthest away from the edge of both the drop and the crystals. This pixel coordinate is used as the suggested coordinate for dispensing compound. Figure 5 shows an example of this process.

**Fig. 5.**
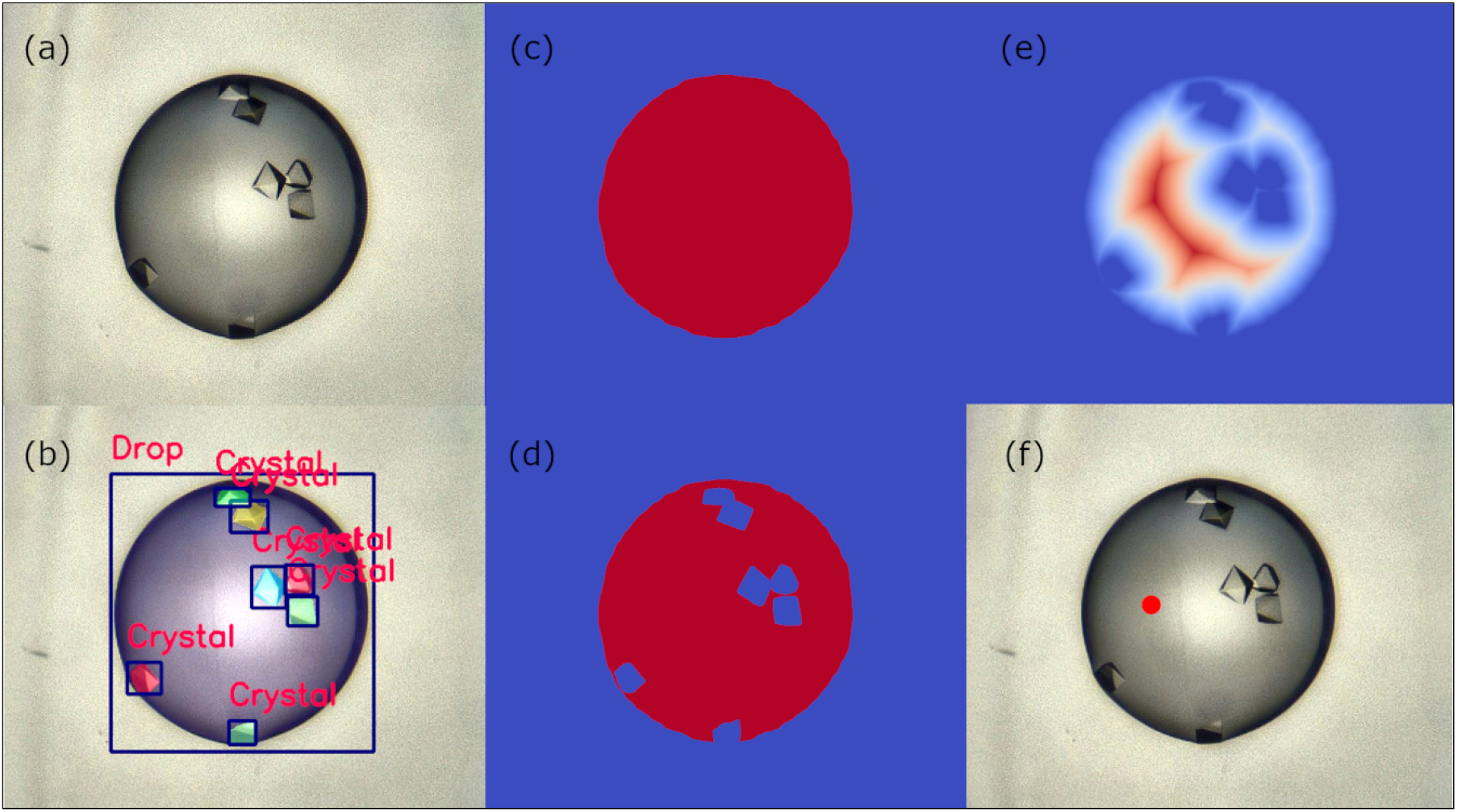
Using the output from the XChem CHiMP Detector Mask-R-CNN model to determine a coordinate to dispense compound for fragment soaking. For the mask images, low pixel values are depicted in blue, high pixel values are depicted in red. (a) Original micrograph of a crystallisation drop. (b) The output bounding boxes, labels and masks from the XChem CHiMP Detector network. Bounding boxes are dark blue, associated object labels are in red. Instance segmentation masks colours are randomised. (c) The predicted mask for the drop. (d) The mask for the drop minus the union of the predicted crystal masks. (e) The Euclidean distance transform of the mask in image (d), red pixels are those farthest from an edge. (f) The original image showing the coordinate determined by taking the maximum pixel value from image (e) as a red dot; this is the suggested dispensing coordinate.

## Results

### 5. Classification Network Performance

After training, the performance of the classification networks was assessed by way of metrics calculated on the validation data as well as on the independent test sets.

#### 5.1. Performance on the VMXi Classification Validation Set

On the validation set of the VMXi Classification Dataset, CHiMP Classifier-v1 achieved a precision, recall and *F*1 score for the *Crystals* class of 0.95, 0.92, and 0.94 respectively. For version 2 of the classification network, the precision, recall and *F*1 metrics for the *Crystals* class were 0.97, 0.88, and 0.93 respectively on this same dataset. The metrics for the other image classes can be found in Supplementary Section 3, Table 11 and Table 12.

#### 5.2. Performance on the VMXi Classification Test Sets

As described in Section 3.5, a set of 1000 images from the robotic imaging microscope on VMXi was collated. All of the images were scored by a panel of three experts and two sets created based upon either unanimous agreement between experts (the unambiguous set) or majority agreement (the mostly unambiguous set). The pairwise agreement between classifications given by the experts ranged from 72.9% to 75.4% showing that the experimental outcomes can be ambiguous to human scorers. All three experts agreed on 63.2% of the images. One of the experts was tasked with scoring the test set of images twice with a period of six months between sessions and was found to agree with themself 83.0% of the time.

The per-class precision, recall and *F*1 metrics for classifying both of these sets of images were calculated for CHiMP Classifier-v1, CHiMP Classifier-v2 and the MARCO Classifier. Table 5 summarises the per-class *F*1 metrics for each model on both of these datasets.

**Table 5.**
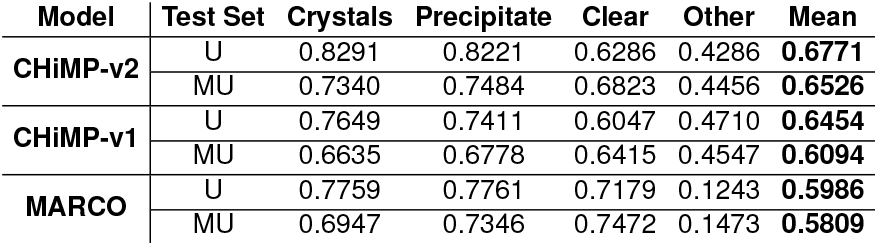
Per-class *F*1 metrics for test sets of images from the VMXi beamline. Test Set U denotes the unambiguous test set of 632 images where three experts agreed on the label. Test Set MU denotes the mostly unambiguous test set of 949 images where two experts agreed on the label.

The precision and recall values for these models can be found in Supplementary Section 3, Table 13, alongside metrics for a number of other models trained using different strategies or with a different architecture for comparison. Other models trained include ConvNeXt Tiny models trained either solely on the MARCO dataset or solely on the VMXi Classification dataset plus a ResNet50 model trained using the same strategy as for the final CHiMP Classifier-v2 ConvNeXt model.

### 6. Detection Network Performance

As described previously (Section 2.1), the most widely metric to describe performance of object detection networks is mean average precision (mAP). This metric was calculated for various IoU thresholds on the validation set of images for the both the VMXi CHiMP Detector and the XChem CHiMP detector networks.

#### 6.1. VMXi CHiMP Detector

The VMXi CHiMP detector network was evaluated on the 47 images in the validation set of the VMXi Detector Dataset and achieved a mAP of 0.31 for bounding boxes on the crystals class when using the standard COCO IoU threshold ranges from 0.5 to 0.95 (with a step size of 0.05). Full mAP results are shown in Table 6. A figure showing eaxmple outputs from the detector network can be seen in Figure 6. A figure showing the display of VMXi CHiMP Detector results in SynchWeb, alongside the plate schematic with a classification overview giving CHiMP Classifier-v2 results, can be seen in Supplementary Section 3, Figure 9 and an additional view of the display of detection results alone can be seen in Figure 6.

**Table 6.**
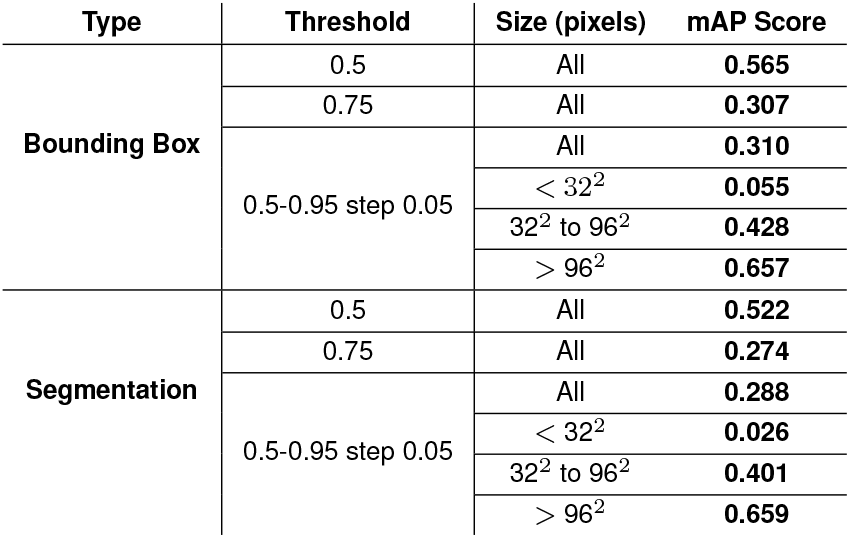
Mean average precision metrics calculated for the VMXi CHiMP Detector Mask-R-CNN for the *Crystals* class. Evaluated on the 47 validation images from the VMXi Detector Dataset

**Fig. 6.**
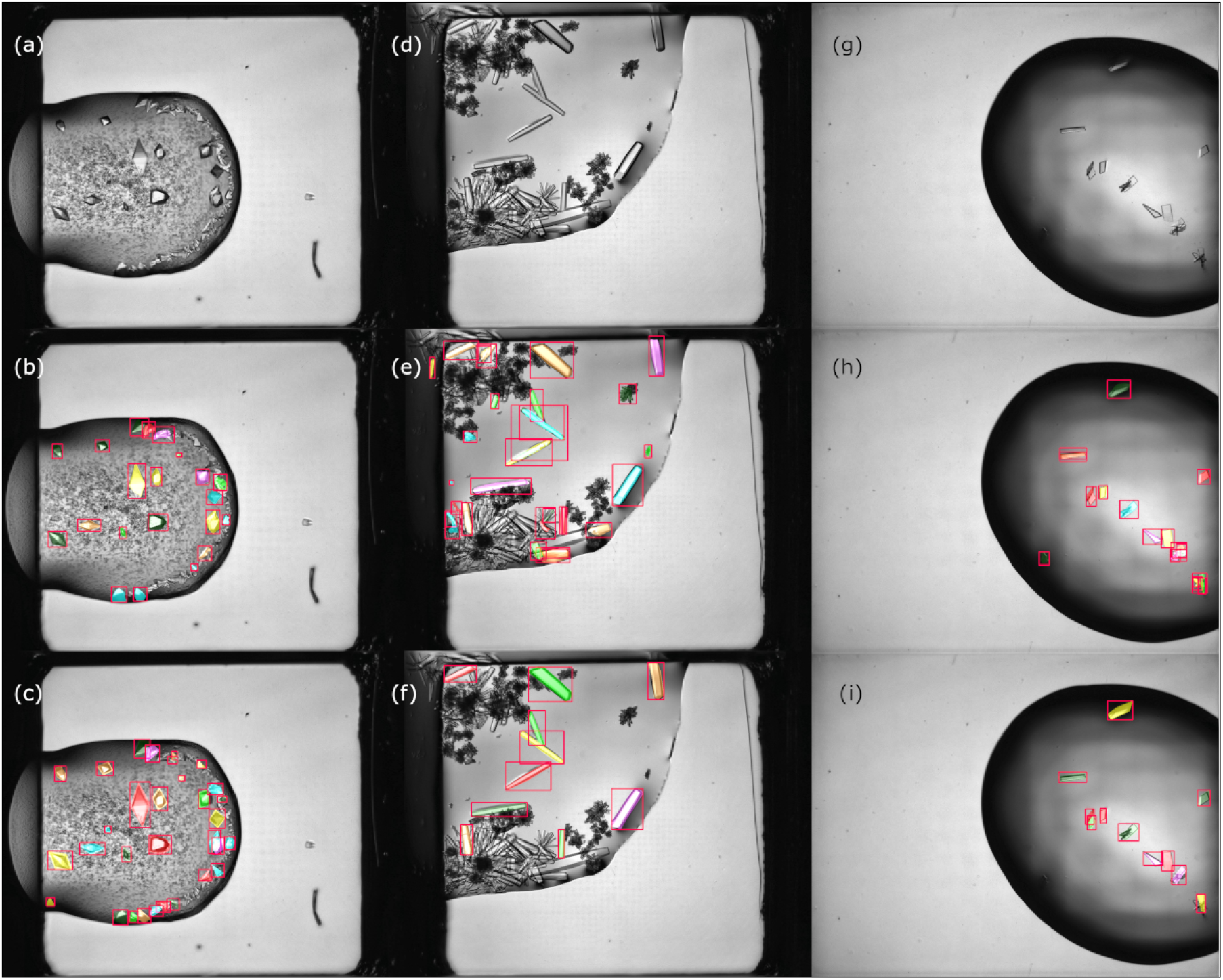
Output from the VMXi CHiMP Detector Mask-R-CNN model on members of the validation set from the VMXi Detector Dataset. Red bounding boxes denote objects in the *Crystals* class. Instance segmentation mask colours are randomised. (a, d, g) Original micrographs of crystallisation drops. (b, e, h) The corresponding bounding boxes and masks as predicted by the VMXi CHiMP Detector network. Objects were predicted with a probability *≥* 0.6. (c, f, i) The ground truth bounding boxes and masks from the VMXi Detector Dataset as annotated by an expert crystallographer.

#### 6.2. XChem CHiMP Detector

The XChem CHiMP detector network was evaluated on the 70 images in the validation set of the VMXi Detector Dataset and achieved a mAP of 0.380 for bounding boxes averaged over the *Crystals* class and the *Drops* class when using the standard COCO IoU threshold ranges from 0.5 to 0.95 (with a step size of 0.05). Full mAP results are shown in Table 7. Example outputs network on members of the validation set can be seen in Figure 7.

**Table 7.**
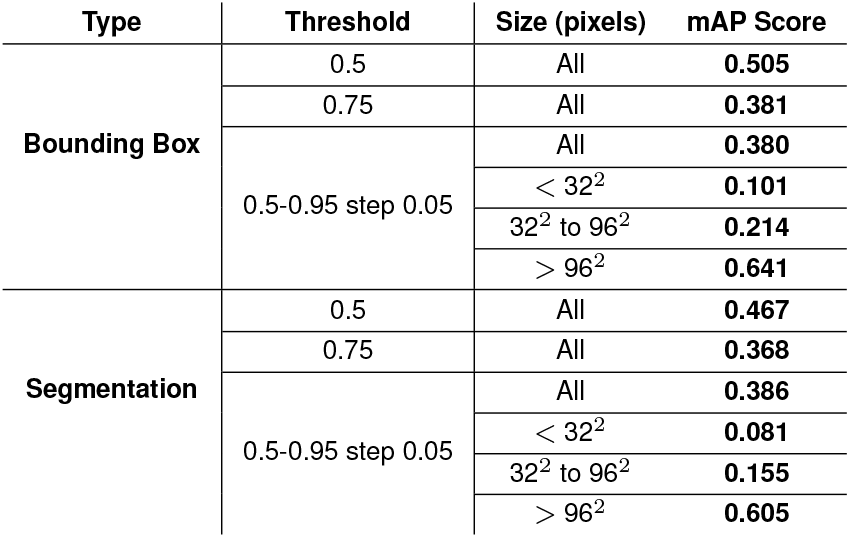
Mean average precision metrics calculated for the XChem CHiMP Detector Mask-R-CNN averaged for the *Crystals* class and the *Drops* class. Evaluated on the 70 validation images in the XChem Detector Dataset

**Table 8.** Summary of label changes during cleaning of the VMXi Classification Dataset. Number Changed is the number of images in that class that have a different label after cleaning to as opposed to before cleaning (rather than the difference in count for the entire category, as given in the Before Cleaning and After Cleaning columns). Percentage Changed is calculated as a percentage of the label count before cleaning.

| Label | Before Cleaning | After Cleaning | Number Changed | Percentage Changed |
| --- | --- | --- | --- | --- |
| Crystals | 8,301 | 8,406 | 249 | 3.0% |
| Clear | 1,514 | 1,529 | 146 | 9.6% |
| Precipitate | 3,037 | 3,101 | 338 | 11.1% |
| Other | 1,099 | 915 | 171 | 15.6% |
| Total | 13,951 | 13,951 | 904 | 6.5% |

**Fig. 7.**
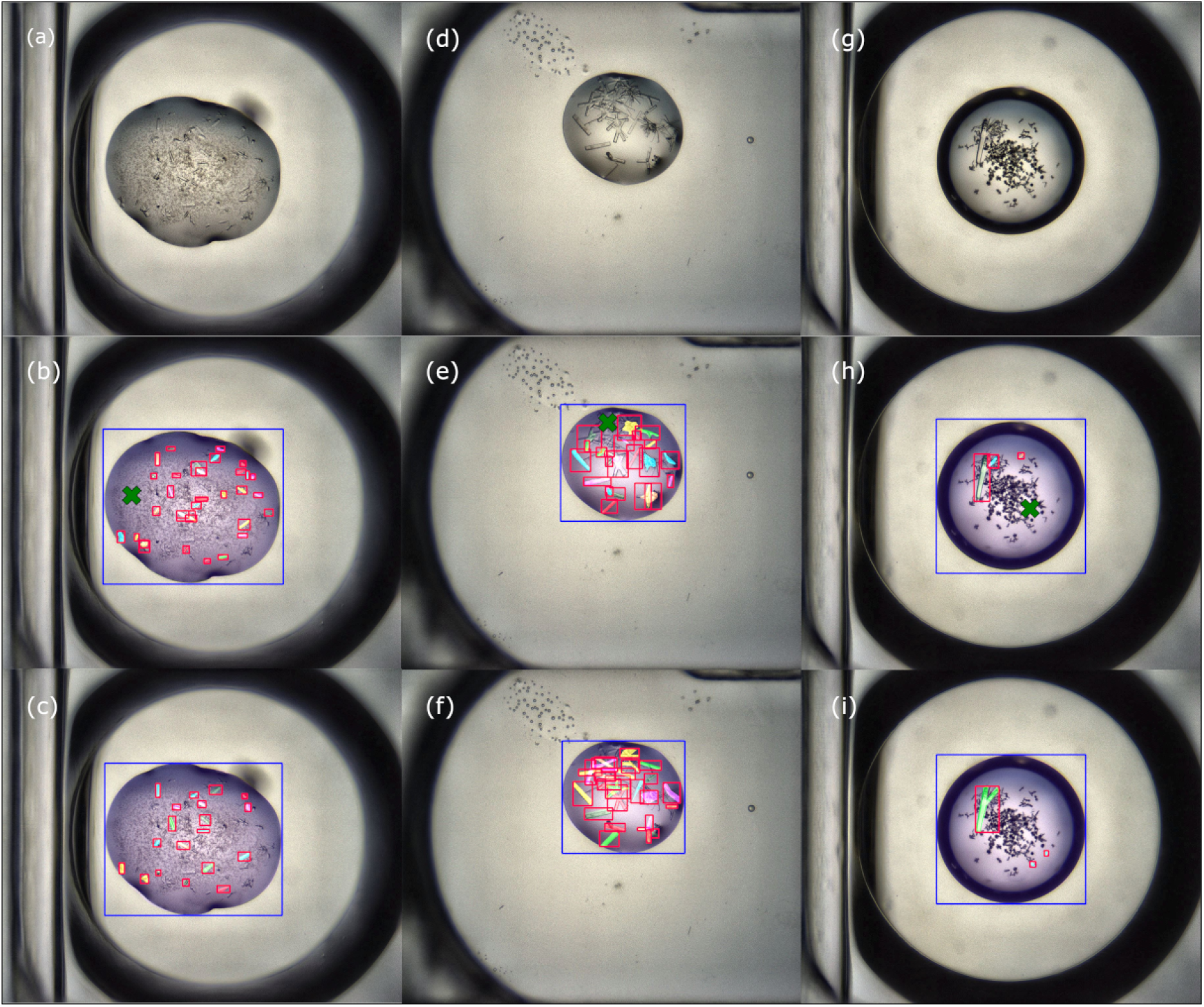
Output from the XChem CHiMP Detector Mask-R-CNN model on members of the validation set from the XChem Detector Dataset. Blue bounding boxes denote objects in the *Drops* class, red bounding boxes denote objects in the *Crystals* class. Instance segmentation mask colours are randomised. Green crosses denote candidate position to dispense compound for fragment soaking (calculated via a distance transform). (a, d, g) Original micrographs of crystallisation drops. (b, e, h) The corresponding bounding boxes and masks as predicted by the XChem CHiMP Detector network. Objects were predicted with a probability *≥* 0.6. Candidate dispensing positions were calculated from output masks. (c, f, i) The ground truth bounding boxes and masks from the XChem Detector Dataset as annotated by an expert crystallographer.

**Fig. 8.**
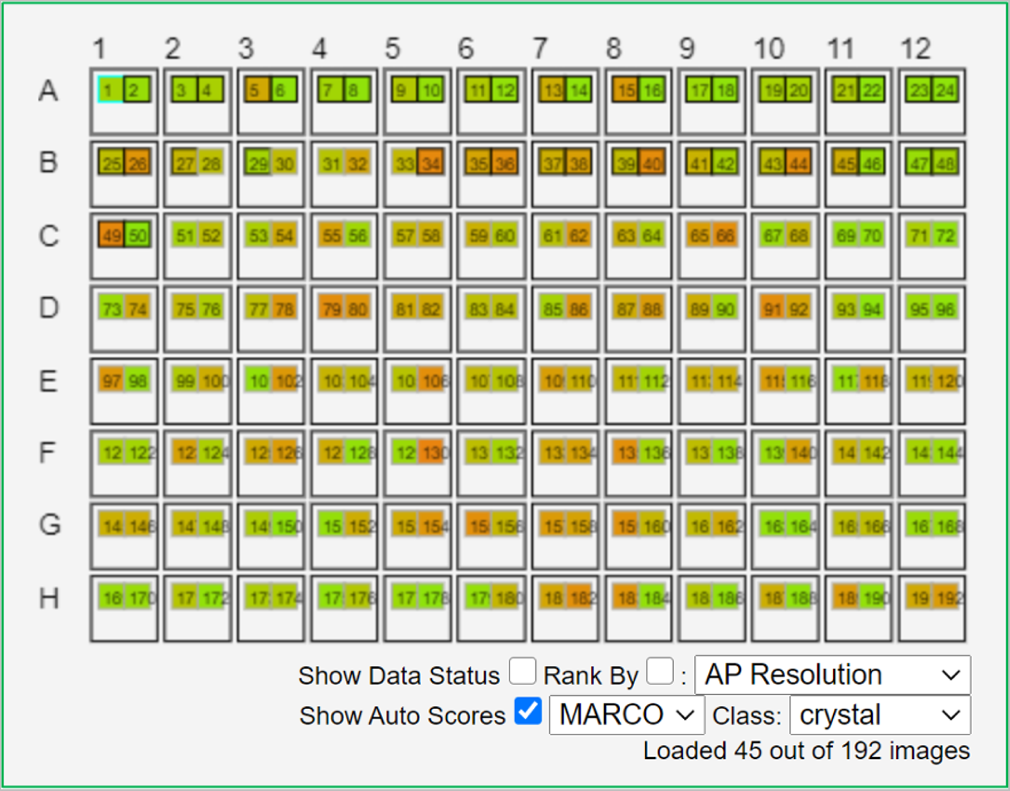
Schematic overview of classification outputs from the MARCO CNN (24), as displayed in SynchWeb (23) for users of the VMXi beamline at Diamond Light Source. The 96 plate wells are represented as larger boxes with the two subwells containing crystallisation droplets shown as smaller boxes within. Each subwell is assigned a colour on a gradient from red to green that represents the probability of the associated image being assigned to the class *Crystals* with green representing a high probability.

**Fig. 9.**
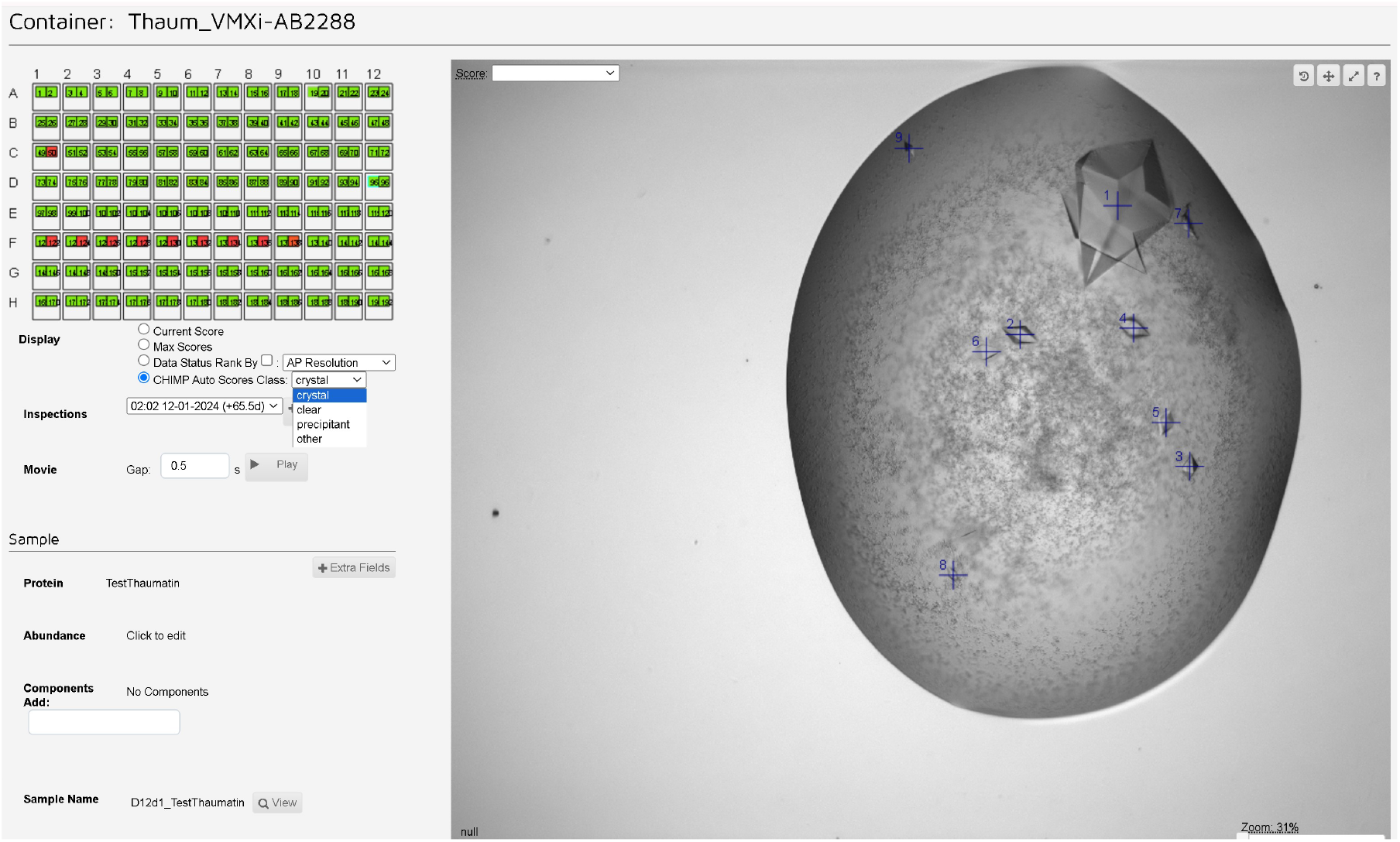
Image of the SynchWeb interface showing a schematic overview of classification outputs from the CHiMP Classifier-v2 CNN alongside an image overlaid with the crystal centroids detected by the VMXi CHiMP Detector network. In the schematic, the 96 plate wells are represented as larger boxes with the two subwells containing crystallisation droplets shown as smaller boxes within. Each subwell is assigned a colour on a gradient from red to green that represents the probability of the associated image being assigned to the class *Crystals* with green representing a high probability. The blue crosses on the associated image mark the crystal centroid positions that can be queued for automated data collection.

**Fig. 10.**
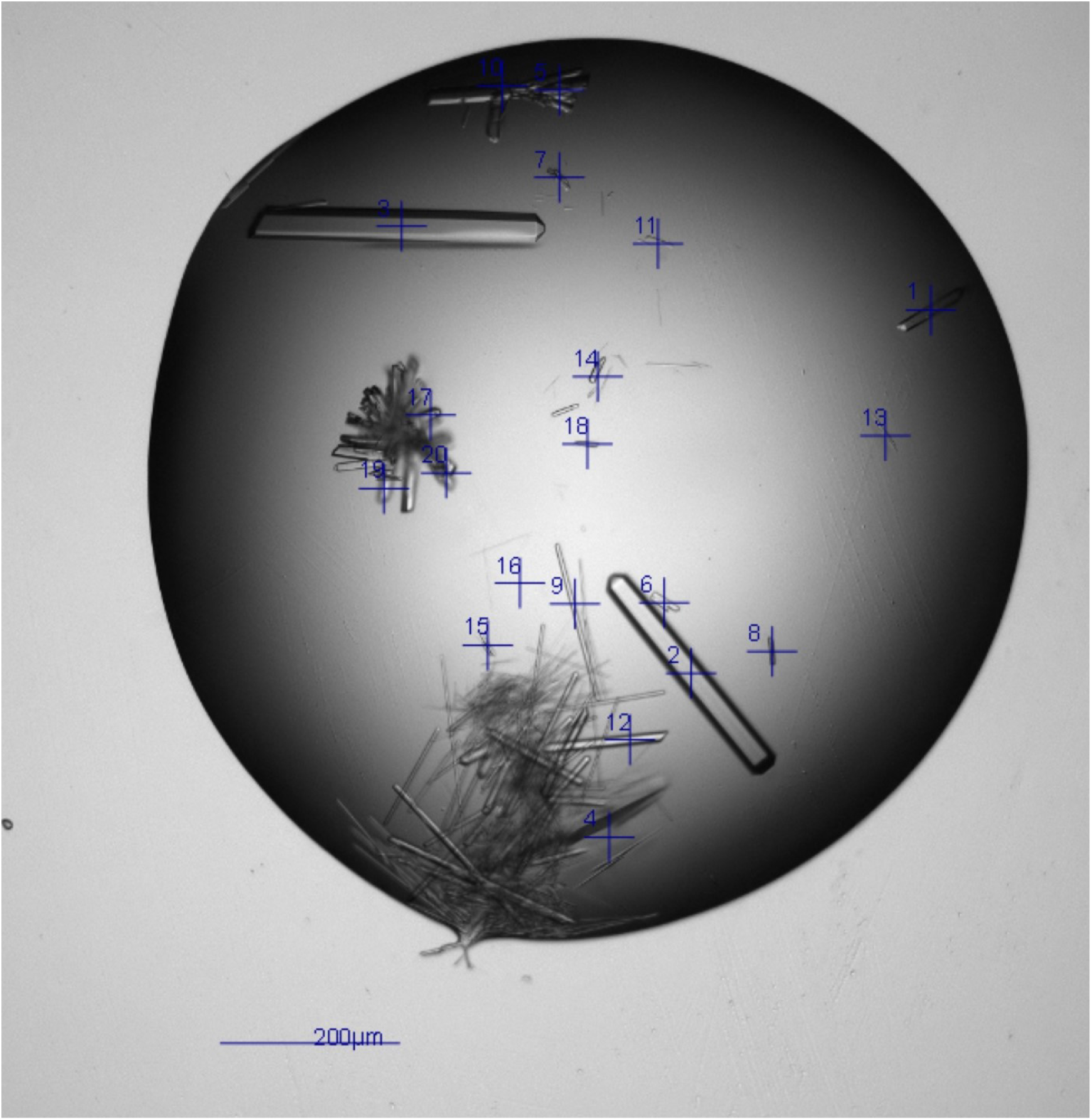
Image from the SynchWeb interface showing an image overlaid with the crystal centroids (blue crosses) detected by the VMXi CHiMP Detector network. The centroid coordinates are calculated from the crystal instance segmentation masks output by the network and are stored in the ISPyB LIMS.

## Discussion

In this study, deep learning networks were fine-tuned, either to classify images into categories of experimental outcome, or to detect and segment objects in the images to allow targeting of locations for data collection or compound dispensing. This was done to deliver increased automation for scientists who normally have to undertake these repetitive and time-consuming tasks.

### 7. On the classification of experimental outcomes

In order to train a network to classify the experimental outcomes, a robust set of data is needed for training. This study has relied on two datasets for image classification, the VMXi Classification Dataset (described in Section 3.1) and the MARCO dataset (24), described in Section 3.2. These datasets both contain labels created by experts but this, in itself, does not mean that these labels are robust. Firstly, as mentioned previously, the categories for classification differ between different institutions, so when images from multiple sources are collated (as in the case of the MARCO dataset) the labels need to be standardised into a system that may not adequately describe the complexity of the outcomes, thereby leading to ambiguity in the categorisation. This oversimplification is compounded by the fact that each image is only given one label when, in fact, an image often contains multiple outcomes. An image that contains both precipitate and a microcrystals will only be given the label *Crystals*. Multilabel classification is a commonly used other fields of study; in satellite remote sensing, for example, where one image may contain forest, savanna and a river, it would be given all three labels. Although studies do exist where a larger number of categories have been used in order to capture the outcome with more granularity, multi-label classification datasets do not exist for protein crystallisation outcomes. This is largely due to historical precedent where the labelling of images is made as simple as possible to ease the burden on annotators. To further ease the burden, annotating images is often optional, which then leads to unbalanced data since the crystallographer is most likely to label crystalline outcomes and to provide no label at all to other outcomes.

The ambiguity between labels assigned by domain experts is highlighted by the process used in this study in order to obtain a “ground truth” test set of images for model evaluation (described in Section 5.2). When categorising 1000 images, three experts agreed only 63% of the time, the maximum pairwise agreement was 75% and the same expert agreed with themself 83% of the time when re-assigning labels on the same images 6 months later. The fact that three experts with many years of combined experience could not agree on a label for 51 of the images also goes to show that the boundaries between categories are flexible. It is this ambiguity that makes the task non-trivial, despite the fact that image classification by CNNs is seen as something that can now be routinely performed with super-human accuracy in other domains (74). The catch-all class of *Other* in particular seems to be ambiguous. When cleaning the labels for the VMXi dataset (described in Supplementary Section 2, Subsection 2) more than 15% of the images in the *Other* category had their label changed by an expert reviewer.

When compared to the MARCO classifier, which has been seen as the most robust model for classifying images of crystallisation droplets in recent times, in this study, we were able to achieve overall higher *F*1 metrics (averaged across all classes) on both our *ambiguous* and *mostly unambiguous* test sets of images from the VMXi beamline. This was achieved by fine-tuning a pre-trained classifier either solely on local images (for CHiMP Classifier-v1) or with a combination of the MARCO dataset and the local images (for CHiMP Classifier-v2), these results are shown in Section 5.2, Table 5. When considering the *Crystals* class alone, however, only the CHiMP Classifier-v2 model (*F*1 0.83 on *unambiguous*, 0.73 on *mostly unambiguous*) was able to surpass the MARCO model (*F*1 0.78 on *unambiguous*, 0.69 on *mostly unambiguous*). Despite this, the effectiveness of the ResNet50 model architecture and training strategy used in the study are highlighted by the fact that the dataset used to train the CHiMP Classifier-v1 model comprised *<* 2.7% of the number of images used to train the MARCO model. An improvement over MARCO for the *Crystals* class was also seen when a ResNet50 model was fine-tuned in the same manner as the CHiMP Classifier-v2 model, achieving an *F*1 metric of 0.82 on the *unambiguous* and 0.71 on the *mostly unambiguous* test sets (Supplementary Section 3, Table 13). This shows that it is not the ConvNeXt architecture alone that is able to outperform the MARCO model. In our study, after a ResNet50 model or a ConvNeXt Tiny model are trained on the MARCO dataset alone they are able to achieve slightly higher average *F*1 metrics than the MARCO classifier. Since the training strategies were different, including using weights from models pre-trained on ImageNet, this improvement cannot be put down to model architecture alone. As a result of the strategy used here, the number of total number of training epochs required to train these models (9 epochs) is much lower than that used to train the MARCO classifier (260 epochs). This shows that using pre-trained weights has the advantage of achieving comparable model performance whilst using fewer computational resources and less energy.

### 8. On the detection of objects in experimental micrographs

Whereas the quality of label data from, sometimes ambiguous, experimental outcomes was often an issue when training classification networks, the quantity of label data was more of an issue when creating a tool to detect and segment instances of objects in the images. The effort involved in creating the inputs required to train a Mask-R-CNN network, namely masks, labels and bounding boxes for every object in the images, were far greater than the single label per image input required for classification. As a result of this, a collaborative approach using the citizen science platform Zooniverse was used (described in Section 4.2). Using this platform was relatively straightforward way to create a web interface that allowed annotation of images by utilising pre-existing drawing tools and allowed multiple annotators to work on the same dataset simultaneously whilst maintaining data integrity. Even with the combined effort of several annotators, however, the size of the resulting datasets was small, with 237 images in the VMXi Detector Dataset and 350 images in the XChem Detector Dataset.

Much like for the classification networks, the limited amount of training data meant that a strategy of fine-tuning a set of pre-trained weights was used, along with use of image augmentations to prevent overfitting to the data. The final mAP metrics (reported in Section 6.1 for the VMXi CHiMP Detector network and Section 6.2 for the XChem CHiMP Detector network) show that the XChem CHiMP detector achieved a higher mAP on both bounding boxes (0.38 vs 0.31) and segmentation (0.39 vs 0.29) for all objects when averaged over the standard IoU thresholds (0.5-0.95 step 0.05). However the mAP performance of the VMXi CHiMP detector network was higher for large objects (*>* 96^2^ pixels) for both bounding boxes (0.66 vs 0.64) and segmentation (0.66 vs 0.61) over the same IoU thresholds than for the XChem Detector network. It is important to bear in mind that the mAP for the XChem Detector is averaged over two classes, however, *Crystals* and *Drops* whereas the VMXi Detector only predict the *Crystals* class.

The complexities of calculating mAP values mean that it is hard to make a direct comparison with the performance of the only other protein crystal detection network where such metrics are reported in the literature. Qin et al. (60) reported a mAP score of 0.70 for their Mask-R-CNN at an IoU threshold of 0.5 for their Mask-R-CNN (presumably for bounding boxes, though they do not specify this) which detects solely the *Crystals* class images from the MARCO dataset. This is higher than the score that was achieved by the VMXi CHiMP Detector network using the same metric (0.57 for bounding boxes and 0.522 for segmentation). The figure reported in the study was calculated on just 10 images and there is no breakdown of performance on different IoU thresholds and different object sizes given, which further complicates a direct comparison.

The utility of both the VMXi CHiMP Detector network and the XChem CHiMP Detector network can be judged by viewing examples of their predicted output on images from their validation sets and comparing the output to the annotation provided by expert crystallographers. Figure 6 and Figure 7 show some of these examples for each of the detector networks. It can be seen that the predicted objects (panels (b), (e) and (h)) in these figures broadly agree with the expert annotation (panels (c),(f) and (i)). There are some instances where crystals annotated by experts are missed (e.g. Figure 6, panels (b) and (c) and Figure 7, panels (h) and (i)), however there are also some instances where the network has annotated crystals that have been missed by the human (e.g Figure 6, panels (e) and (f) plus panels (h) and (i)). For the XChem CHiMP detector, using the predicted drop and crystal masks to calculate a candidate position for dispensing compound appears to provide coordinates that are in a location away from the bulk of mountable crystals an also away from the edge of the drop.

## Conclusions

The ultimate value of the automatic classification of images into categories of experimental outcome is realised when the model is integrated into the software interface that scientists use to monitor their experiments. In this case, a schematic overview of the subwells in the crystallisation plate in the VMXi SynchWeb interface allows the locations of various outcomes to be quickly found along with a general overview of the proportion of experiments that resulted in each outcome. This overview of outcomes can then be used as the entry point to viewing images that are likely to be of interest and that are further annotated by an object detection network to pinpoint any crystals that are present. An example of this interface can be seen in Supplementary Section 3, Figure The fact that the classification tool created in this study has been shown to achieve performance metrics that surpass those achieved by the MARCO tool (itself seen as one that could be trusted by crystallographers weary of missing important results), hopefully means that many hours otherwise spent performing manual inspection of images will be saved by this work. Integrating the VMXi CHiMP Detector tool into the SynchWeb interface has meant that human interaction is no longer required to mark the location of the crystals and add them to a queue for X-ray data collection. This, in turn, means that the experimental station at the beamline has the potential to work fully autonomously, with setting up the experiments in the crystal plate and adding the information to the ISPyB database being the only steps that require human intervention. After this is done, plate inspections, data collection and protein structure determination can now happen fully automatically.

By using the XChem CHiMP detector network, candidate positions for the dispensing of small molecule fragment solutions into the droplets can now be calculated automatically, thereby speeding up the process of setting up soaking experiments during screening campaigns, where hundreds of such compounds are soaked into crystals prior to X-ray data collection.

It is hoped that sharing the training data, as well as the model weights and the code for model training and prediction of the networks described here will aid other researchers to take advantage of these tools, as well as to serve as a starting point for further improvements. It is also hoped that this work will ultimately lead to faster structural discoveries that will have the potential to further our understanding of disease processes and how to treat them.

## Data availability

### VMXi Classification Dataset

The 13,951 images that make up this dataset and the 1000 images that form the test sets of images used to evaluate model performance are available at doi:10.5281/zenodo.11097395.

### VMXi Detector & XChem Detector Datasets

The 237 images and 350 images that make up the VMXi CHiMP Detector and XChem CHiMP Detector Datasets respectively, along with the corresponding image masks for drops and crystals, are available at doi:10.5281/zenodo.11110372.

### CHiMP Classifier-v2 Network

The model weights for the ConvNeXt-Tiny CHiMP Classification-v2 Network are available at doi:10.5281/zenodo.11190973.

### VMXi CHiMP Detector Network

The model weights for the Mask-R-CNN VMXi CHiMP Detector Network are available at doi:10.5281/zenodo.11164787.

### XChem CHiMP Detector Network

The model weights for the Mask-R-CNN XChem CHiMP Detector Network are available at doi:10.5281/zenodo.11165194.

### CHiMP Tools

The software repository containing methods used for training and inference of the classification and detection networks discussed in this study is available at doi:10.5281/zenodo.11244711.

## ACKNOWLEDGEMENTS

We would like to thank the VMXi beamline staff for helpful discussions and for time spent annotating images, particularly Juan Sanchez-Weatherby. We would like to thank Urszula Neuman, Stuart Fisher, Victor Nwaiwu and Emma Dixon for their work on integrating the classification and detection networks into the beamline software pipeline and also Richard Gildea and Graeme Winter for discussions on how to do so. We would also like to thank the XChem staff who spent time annotating images to produce training data for the object detection network, namely: Daren Fearon, Charlie Tomlinson, Peter Marples, Isabel Barker and Alexandre Dias.

## Supplementary Section 1: Initial Installation and Evaluation of MARCO at the DLS VMXi Beam-line

### 1. Initial assessment of MARCO performance on images from the VMXi facility

To evaluate the performance of the MARCO model on micrographs captured in-house, the images from two crystal plates, each with 288 subwells was scored by an expert using the same four categories used by MARCO: *Crystals, Precipitate, Clear* and *Other*. One plate was from an experiment set-up to crystallise haemoglobin and the other contained thermolysin. From the 576 classified images, 20 were selected at random from each of the four classes giving a final, balanced, dataset of 80 images. These were then scored using MARCO for comparison, giving a precision, recall and *F*1 score for the *Crystals* class of 0.82, 0.7 and 0.76 respectively. The recall and *F*1 metric scores are moderately lower than those found by Bruno et al. (24) for their independent test set of images (precision 0.78, a recall of 0.87 and *F*1 score of 0.82) but the precision is comparable. It was found that MARCO also tended to over-classify images as Precipitate, with 22 false positive classifications. This resulted in a recall of 100% of the *Precipitate* images but this was at the expense of precision which was 0.48 (*F*1 metric 0.65).

## Supplementary Section 2: Appendix to Methods

### 1. Details on the collection of images at the VMXi beamline experimental facility

All experimental crystal plates on the VMXi beamline are registered as *containers* in the ISPyB LIMS relational database before being inserted into a Rock Imager 1000 (Formulatrix, USA) automated microplate imager. Images of every crystallisation sub-well are recorded (termed an *inspection*) immediately after insertion of the experimental plate and, in addition, subsequent inspections are carried out on a Fibonacci sequence schedule (i.e. 1, 3, 5, 8 … days) to monitor the experiments over time. A *z*-stack of images, taken with different focal points within the plate subwell, is combined into an extended focus image and then saved as JPEG format with a resolution of 3376 × 2704 pixels. Once created, all images are moved to directories linked to the experimental visit that the container is part of and also to the inspection number. This information is also recorded in the ISPyB LIMS (21). Once recorded, the images are available to view in the SynchWeb browser interface (23). In this interface, scientists are able to browse the images and provide a score for classifying the content of the image.

### 2. Cleaning of the data labels for the VMXi Classification Dataset

A ResNet50 model initialised with ImageNet weights was fine-tuned with a random split of 80% of images in the training set and 20% of images in the validation set. The *Image-Cleaner* functionality of the *fastai* library (65) was then used to correct mislabelled images. This process entails using the trained ResNet50 to classify the entire dataset of images (training and validation set) whilst also recording the cross entropy loss for the classifications. Since a high loss value for a classification can be equated with a high uncertainty, images can then viewed in order of the most uncertain label first. The original label given in the dataset is displayed alongside the image and the label can then be corrected if it is wrong; this is more time-efficient than reviewing the entire dataset.

### 3. Training the CHiMP Classifier-V1 ResNet50 CNN

At the start of each phase of model training, the weights of the ResNet50 convolutional layers were “frozen”, i.e. gradients were not calculated and the weights were not updated, only the weights of the fully connected classification head network, responsible for determining the posterior probability of the four image classes were available to be modified. After five epochs of training on the partially frozen model, all of the models were “unfrozen” allowing them to be updated before the next round of model training.

The learning rate to be used was determined through a process of training the model for one epoch whilst exponentially increasing the learning rate for each batch and plotting a graph of loss against learning rate. The rate is chosen from this graph according to the loss profile and this value is then used to define a range of values to be used in a “1Cycle” training strategy (69). In this strategy, which has been shown to accelerate model convergence, the learning rate is gradually increased to a maximum value during the middle of the training epochs and reduced to a low value towards the end. Model parameter optimisation was carried out using AdamW (68).

In the first phase of training, an image size of 128 × 128 pixels was used followed by 256 × 256 pixels in the second phase and 512 × 512 pixels for the final phase. This strategy is summarised in Table 9.

**Table 9.**
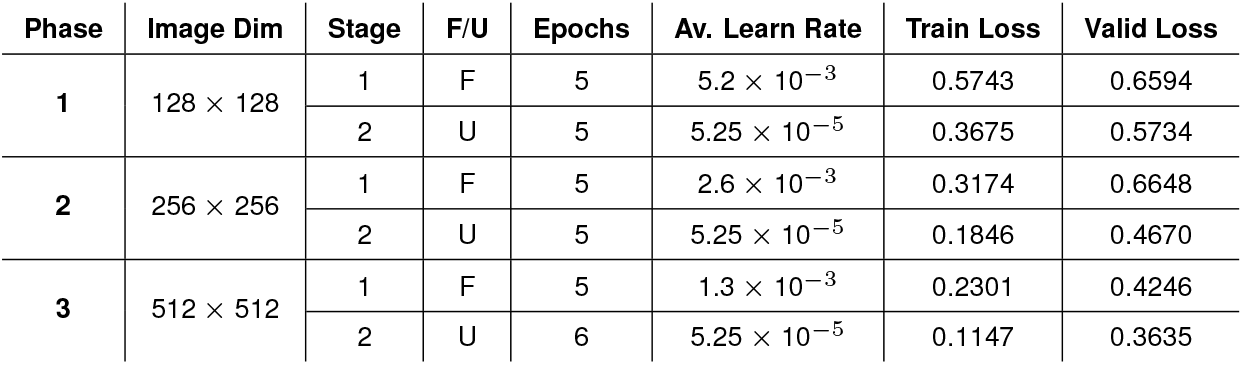
Summary of the training phases for the CHiMP Classifier-v1 ResNet50 CNN using the VMXi Classification Dataset. Image dimensions in pixels, F/U refers to Frozen or Unfrozen model weights respectively. Av Learn Rate refers to average learning rate used during a cyclic learning rate strategy. Loss is cross entropy loss.

**Table 10.** Summary of the training phases for the CHiMP Classifier-v2 ConvNeXt Tiny CNN using the MARCO dataset (M) and then the VMXi Classification Dataset (V). Image dimensions in pixels, F/U refers to Frozen or Unfrozen model weights respectively. Av Learn Rate refers to average learning rate used during a cyclic learning rate strategy. Loss is cross entropy loss.

| Phase | Image Dim | Data | F/U | Epochs | Av. Learn Rate | Train Loss | Valid Loss |
| --- | --- | --- | --- | --- | --- | --- | --- |
| 1 | $128 \times 128$ | M | F | 1 | $3.9 \times 10^{-4}$ | 0.9586 | 0.7380 |
| | | M | U | 3 | $2.6 \times 10^{-5}$ | 0.3722 | 0.3026 |
| 2 | $256 \times 256$ | M | U | 4 | $2.6 \times 10^{-5}$ | 0.1939 | 0.2422 |
| 3 | $512 \times 512$ | M | U | 4 | $2.6 \times 10^{-5}$ | 0.2013 | 0.2136 |
| 4 | $256 \times 256$ | V | F | 1 | $2.6 \times 10^{-5}$ | 1.5590 | 1.7364 |
| | | V | U | 9 | $2.6 \times 10^{-5}$ | 0.6810 | 0.6634 |
| 5 | $512 \times 512$ | V | U | 14 | $2.6 \times 10^{-5}$ | 0.4206 | 0.5220 |

**Table 11.** Classification Performance of CHiMP Classifier-v1 on validation images from the VMXi Classification Dataset

| Class | Precision | Recall | F1 |
| --- | --- | --- | --- |
| Crystals | 0.9525 | 0.9208 | 0.9367 |
| Clear | 0.8392 | 0.9255 | 0.8803 |
| Precipitate | 0.8431 | 0.8569 | 0.8500 |
| Other | 0.5856 | 0.6339 | 0.6089 |

**Table 12.** Classification Performance of CHiMP Classifier-v2 on validation images from the VMXi Classification Dataset

| Class | Precision | Recall | F1 |
| --- | --- | --- | --- |
| Crystals | 0.9701 | 0.8839 | 0.9250 |
| Clear | 0.8457 | 0.9326 | 0.8870 |
| Precipitate | 0.8560 | 0.8331 | 0.8444 |
| Other | 0.5297 | 0.9235 | 0.6733 |

**Table 13.** Per-class precision, recall and *F*1 metrics for classification models for test sets of images from the VMXi beamline. Test Set U denotes the unambiguous test set of 632 images where three experts agreed on the label. Test Set MU denotes the mostly unambiguous test set of 949 images where two experts agreed on the label. Models **CHiMP Classifier-v1, CHiMP Classifier-v2** and **MARCO** as described in main text. **ResNet50-MV** model trained on both MARCO and VMXi classification datasets in the same fashion as CHiMP-v2. **ResNet50-M** model trained solely on MARCO Classification Dataset (prior to additional training on the VMXi Classification Dataset to create ResNet50-MV). **ConvNeXt-M** model trained solely on MARCO Classification Dataset (prior to additional training on the VMXi Classification Dataset to create CHiMP Classifier-v2). **ConvNeXt-V** model trained solely on the VMXi Classification Dataset.

| Model | T-Set | Crystals |  |  | Precipitate |  |  | Clear |  |  | Other |  |  | Average |  |  |
| --- | --- | --- | --- | --- | --- | --- | --- | --- | --- | --- | --- | --- | --- | --- | --- | --- |
|  |  | Prec | Rec | F1 | Prec | Rec | F1 | Prec | Rec | F1 | Prec | Rec | F1 | Prec | Rec | F1 |
| CHiMP-v2 | U | 0.7661 | 0.9034 | 0.8291 | 0.9194 | 0.7435 | 0.8221 | 0.4757 | 0.9263 | 0.6286 | 0.6000 | 0.3333 | 0.4286 | **0.6903** | **0.7266** | **0.6771** |
|  | MU | 0.6564 | 0.8324 | 0.7340 | 0.8640 | 0.6601 | 0.7484 | 0.5575 | 0.8791 | 0.6823 | 0.5399 | 0.3793 | 0.4456 | **0.6544** | **0.6877** | **0.6526** |
| ResNet50-MV | U | 0.7651 | 0.8759 | 0.8167 | 0.8557 | 0.7478 | 0.7981 | 0.4886 | 0.9053 | 0.6347 | 0.5393 | 0.2963 | 0.3825 | **0.6622** | **0.7063** | **0.6580** |
|  | MU | 0.6495 | 0.7765 | 0.7074 | 0.8182 | 0.6573 | 0.7290 | 0.5477 | 0.8516 | 0.6667 | 0.4759 | 0.3405 | 0.3970 | **0.6228** | **0.6565** | **0.6250** |
| CHiMP-v1 | U | 0.7011 | 0.8414 | 0.7649 | 0.8902 | 0.6348 | 0.7411 | 0.4785 | 0.8211 | 0.6047 | 0.5267 | 0.4259 | 0.4710 | **0.6492** | **0.6808** | **0.6454** |
|  | MU | 0.5761 | 0.7821 | 0.6635 | 0.8354 | 0.5702 | 0.6778 | 0.5620 | 0.7473 | 0.6415 | 0.4661 | 0.4440 | 0.4547 | **0.6099** | **0.6359** | **0.6094** |
| ResNet50-M | U | 0.6000 | 0.7862 | 0.6806 | 0.8316 | 0.6870 | 0.7524 | 0.5111 | 0.9684 | 0.6691 | 0.5972 | 0.2654 | 0.3675 | **0.6350** | **0.6768** | **0.6174** |
|  | MU | 0.5316 | 0.7039 | 0.6058 | 0.7860 | 0.5983 | 0.6794 | 0.5475 | 0.9505 | 0.6948 | 0.5440 | 0.2931 | 0.3810 | **0.6023** | **0.6365** | **0.5902** |
| ConvNeXt-M | U | 0.5043 | 0.8069 | 0.6207 | 0.8660 | 0.7304 | 0.7925 | 0.5506 | 0.9158 | 0.6877 | 0.6875 | 0.2037 | 0.3143 | **0.6521** | **0.6642** | **0.6038** |
|  | MU | 0.4596 | 0.7318 | 0.5647 | 0.7930 | 0.6348 | 0.7051 | 0.5700 | 0.9176 | 0.7032 | 0.6163 | 0.2284 | 0.3333 | **0.6097** | **0.6282** | **0.5766** |
| MARCO | U | 0.7532 | 0.8000 | 0.7759 | 0.6636 | 0.9348 | 0.7762 | 0.6043 | 0.8842 | 0.7179 | 0.7333 | 0.0679 | 0.1243 | **0.6886** | **0.6717** | **0.5986** |
|  | MU | 0.6966 | 0.6927 | 0.6947 | 0.6352 | 0.8708 | 0.7346 | 0.6381 | 0.9011 | 0.7472 | 0.7308 | 0.0819 | 0.1473 | **0.6752** | **0.6366** | **0.5809** |
| ConvNeXt-V | U | 0.5789 | 0.8345 | 0.6836 | 0.8667 | 0.5652 | 0.6842 | 0.4940 | 0.8737 | 0.6312 | 0.4381 | 0.2840 | 0.3446 | **0.5944** | **0.6393** | **0.5859** |
|  | MU | 0.5000 | 0.7765 | 0.6083 | 0.7932 | 0.5281 | 0.6341 | 0.5659 | 0.8022 | 0.6636 | 0.4148 | 0.3147 | 0.3578 | **0.5685** | **0.6054** | **0.5660** |

### 4. Training the CHiMP Classifier-V2 ConvNeXt Tiny CNN

Initial fine-tuning of the model on the MARCO Dataset was done for 12 epochs in total, split across three phases with increasing image dimensions at each phase (starting with 128 × 128 pixels, then 256 × 256 pixels, then 512 × 512 pixels). Training and validation sets of data were the same as those publicly released by Bruno et al. (24) and used to train their MARCO classification model (a breakdown of image classes can be seen in Table 3). The batch size used was 24. Due to the relatively large size of this dataset, training was distributed across four NVIDA Tesla V100S GPUs, each equipped with 32 GB of VRAM. The cross entropy loss for the training set was 0.201 and for the validation set was 0.214 after this section of training process.

This model was then fine-tuned further on the VMXi Classification dataset (described in Section 3.1). During this process, the model was found to have a propensity to over-fit on this smaller dataset, with the training loss dropping much lower than the validation loss. As a result, more regularisation was added in the form of stochastic depth (75) with a probability of 0.2. In this technique, layers of the network are randomly dropped during training, shortening the network and forcing the model to adapt to a situation where not all layers are used, this allows the model to be trained for longer without over-fitting. A two-phase strategy was used with first phase using images with dimensions of 256 × 256 pixels for 10 epochs. The model weights were frozen for the first epoch, training only the network head. In the second phase, using images of 512 × 512 pixels, the model was fine-tuned for 14 epochs. The final training loss was 0.421 and the final validation loss was 0.522.

## Supplementary Section 3: Appendix to Results

### 1. Performance of networks on the VMXi Classification Dataset validation set

Tables 11 and 12 show the per-class metrics for CHiMP Classifier-v1 and CHiMP Classifier-v2 on the validation images from the VMXi Classification Dataset.

### 2. Classification Performance on Test Sets of Images from the VMXi Beamline

Performance metrics for a number of different classification models on the test sets of images (described in Section 3.5) are shown here.

### 3. SynchWeb visualisation of the output of the classification and detection networks for users of the VMXi experimental facility

The outputs from the CHiMP Classifier-v2 network and the VMXi CHiMP detector network are stored in the ISPyB LIMS. Using the SynchWeb web interface, facility users can browse this information in the form of a plate overview schematic alongside images with overlaid crystal centroids that are displayed by clicking on a subwell.

## Notes

https://zenodo.org/doi/10.5281/zenodo.11097395

https://zenodo.org/doi/10.5281/zenodo.11110372

https://zenodo.org/doi/10.5281/zenodo.11190973

https://zenodo.org/doi/10.5281/zenodo.11164787

https://zenodo.org/doi/10.5281/zenodo.11165194

https://zenodo.org/doi/10.5281/zenodo.11244711

### Competing Interest Statement

The authors have declared no competing interest.

## Bibliography

1. John Jumper, Richard Evans, Alexander Pritzel, Tim Green, Michael Figurnov, Olaf Ronneberger, Kathryn Tunyasuvunakool, Russ Bates, Augustin Žídek, Anna Potapenko, Alex Bridgland, Clemens Meyer, Simon A. A. Kohl, Andrew J. Ballard, Andrew Cowie, Bernardino Romera-Paredes, Stanislav Nikolov, Rishub Jain, Jonas Adler, Trevor Back, Stig Petersen, David Reiman, Ellen Clancy, Michal Zielinski, Martin Steinegger, Michalina Pacholska, Tamas Berghammer, Sebastian Bodenstein, David Silver, Oriol Vinyals, Andrew W. Senior, Koray Kavukcuoglu, Pushmeet Kohli, and Demis Hassabis. Highly accurate protein structure prediction with AlphaFold. Nature, 596(7873):583–589, August 2021. doi:10.1038/s41586-021-03819-2.

2. Yifan Cheng. Single-particle cryo-EM—How did it get here and where will it go. Science, 361(6405):876–880, August 2018. doi:10.1126/science.aat4346.

3. J. T. Ng, C. Dekker, P. Reardon, and F. von Delft. Lessons from ten years of crystallization experiments at the SGC. Acta Crystallographica Section D: Structural Biology, 72(2):224–235, February 2016. doi:10.1107/S2059798315024687.

4. J. Jancarik and S.-H. Kim. Sparse matrix sampling: A screening method for crystallization of proteins. Journal of Applied Crystallography, 24(4):409–411, August 1991. doi:10.1107/S0021889891004430.

5. Alice Douangamath, Ailsa Powell, Daren Fearon, Patrick M. Collins, Romain Talon, Tobias Krojer, Rachael Skyner, Jose Brandao-Neto, Louise Dunnett, Alexandre Dias, Anthony Aimon, Nicholas M. Pearce, Conor Wild, Tyler Gorrie-Stone, and Frank von Delft. Achieving Efficient Fragment Screening at XChem Facility at Diamond Light Source. JoVE (Journal of Visualized Experiments), (171):e62414, May 2021. doi:10.3791/62414.

6. T. Moreno-Chicano, L. M. Carey, D. Axford, J. H. Beale, R. B. Doak, H. M. E. Duyvesteyn Ebrahim, R. W. Henning, D. C. F. Monteiro, D. A. Myles, S. Owada, D. A. Sherrell, M. L. Straw, V. Šrajer, H. Sugimoto, K. Tono, T. Tosha, I. Tews, M. Trebbin, R. W. Strange, K. L. Weiss, J. a. R. Worrall, F. Meilleur, R. L. Owen, R. A. Ghiladi, and M. A. Hough. Complementarity of neutron, XFEL and synchrotron crystallography for defining the structures of metalloenzymes at room temperature. IUCrJ, 9(5):610–624, September 2022. doi:10.1107/S2052252522006418.

7. Marcus Fischer. Macromolecular room temperature crystallography. Quarterly Reviews of Biophysics, 54:e1, January 2021. ISSN 0033-5835, 1469-8994. doi:10.1017/S0033583520000128.

8. J. H. Beale, R. Bolton, S. A. Marshall, E. V. Beale, S. B. Carr, A. Ebrahim, T. Moreno-Chicano, M. A. Hough, J. a. R. Worrall, I. Tews, and R. L. Owen. Successful sample preparation for serial crystallography experiments. Journal of Applied Crystallography, 52(6): 1385–1396, December 2019. doi:10.1107/S1600576719013517.

9. R. E. Thorne. Determining biomolecular structures near room temperature using x-ray crystallography: concepts, methods and future optimization. 79(1):78–94. ISSN 2059-7983. doi:10.1107/S2059798322011652.

10. Katherine E. McAuley, Mark Williams, and Stuart Fisher. BART – the New Robotic Sample Changer for MX Beamlines at Diamond -- Diamond Light Source. Technical report, Diamond Light Source, 2015.

11. F. Cipriani, F. Felisaz, L. Launer, J.-S. Aksoy, H. Caserotto, S. Cusack, M. Dallery, F. diChiaro, M. Guijarro, J. Huet, S. Larsen, M. Lentini, J. McCarthy, S. McSweeney, R. Ravelli, M. Renier, C. Taffut, A. Thompson, G. A. Leonard, and M. A. Walsh. Automation of sample mounting for macromolecular crystallography. Acta Crystallographica Section D: Biological Crystallography, 62(10):1251–1259, October 2006. doi:10.1107/S0907444906030587.

12. E. O. Lazo, S. Antonelli, J. Aishima, H. J. Bernstein, D. Bhogadi, M. R. Fuchs, N. Guichard, S. McSweeney, S. Myers, K. Qian, D. Schneider, G. Shea-McCarthy, J. Skinner, R. Sweet, L. Yang, and J. Jakoncic. Robotic sample changers for macromolecular X-ray crystallography and biological small-angle X-ray scattering at the National Synchrotron Light Source II. Corrigendum. Journal of Synchrotron Radiation, 29(1):280–280, January 2022. doi:10.1107/S1600577521013205.

13. Stephen R. Wasserman, Jordi Benach, John W. Koss, and Laura L. Morisco. The Evolution of High-Throughput Macromolecular Crystallography at Synchrotrons. Synchrotron Radiation News, 28(6):4–9, November 2015. doi:10.1080/08940886.2015.1101320.

14. J. Song, D. Mathew, S. A. Jacob, L. Corbett, P. Moorhead, and S. M. Soltis. Diffraction-based automated crystal centering. Journal of Synchrotron Radiation, 14(2):191–195, March 2007. doi:10.1107/S0909049507004803.

15. E. Pohl, U. Ristau, T. Gehrmann, D. Jahn, B. Robrahn, D. Malthan, H. Dobler, and C. Hermes. Automation of the EMBL Hamburg protein crystallography beamline BW7B. Journal of Synchrotron Radiation, 11(5):372–377, September 2004. doi:10.1107/S090904950401516X.

16. S. Ito, G. Ueno, and M. Yamamoto. DeepCentering: Fully automated crystal centering using deep learning for macromolecular crystallography. Journal of Synchrotron Radiation, 26(4): 1361–1366, July 2019. doi:10.1107/S160057751900434X.

17. Jonathan Schurmann, Isaak Lindhè, Jorn W. Janneck, Gustavo Lima, and Zdenek Matej. Crystal centering using deep learning in X-ray crystallography. In 2019 53rd Asilomar Conference on Signals, Systems, and Computers, pages 978–983, November 2019. doi:10.1109/IEEECONF44664.2019.9048793.

18. J. Sanchez-Weatherby, J. Sandy, H. Mikolajek, C. M. C. Lobley, M. Mazzorana, J. Kelly, G. Preece, R. Littlewood, and T. L.-M. Sørensen. VMXi: A fully automated, fully remote, high-flux in situ macromolecular crystallography beamline. Journal of Synchrotron Radiation, 26(1):291–301, January 2019. doi:10.1107/S1600577518015114.

19. Z. Ren, C. Wang, H. Shin, S. Bandara, I. Kumarapperuma, M. Y. Ren, W. Kang, and X. Yang. An automated platform for in situ serial crystallography at room temperature. IUCrJ, 7(6): 1009–1018, November 2020. doi:10.1107/S2052252520011288.

20. Robert D. Healey, Shibom Basu, Anne-Sophie Humm, Cedric Leyrat, Xiaojing Cong, Jérôme Golebiowski, Florine Dupeux, Andrea Pica, Sébastien Granier, and José Antonio Márquez. An automated platform for structural analysis of membrane proteins through serial crystallography. Cell Reports Methods, 1(6):100102, October 2021. doi:10.1016/j.crmeth.2021.100102.

21. Solange Delagenière, Patrice Brenchereau, Ludovic Launer, Alun W. Ashton, Ricardo Leal, Stéphanie Veyrier, José Gabadinho, Elspeth J. Gordon, Samuel D. Jones, Karl Erik Levik, Seán M. McSweeney, Stéphanie Monaco, Max Nanao, Darren Spruce, Olof Svensson, Martin A. Walsh, and Gordon A. Leonard. ISPyB: An information management system for synchrotron macromolecular crystallography. Bioinformatics, 27(22):3186–3192, November 2011. doi:10.1093/bioinformatics/btr535.

22. R. J. Gildea, J. Beilsten-Edmands, D. Axford, S. Horrell, P. Aller, J. Sandy, J. Sanchez-Weatherby, C. D. Owen, P. Lukacik, C. Strain-Damerell, R. L. Owen, M. A. Walsh, and G. Winter. Xia2.multiplex: A multi-crystal data-analysis pipeline. Acta Crystallographica Section D: Structural Biology, 78(6):752–769, June 2022. doi:10.1107/S2059798322004399.

23. S. J. Fisher, K. E. Levik, M. A. Williams, A. W. Ashton, and K. E. McAuley. SynchWeb: A modern interface for ISPyB. Journal of Applied Crystallography, 48(3):927–932, June 2015. doi:10.1107/S1600576715004847.

24. Andrew E. Bruno, Patrick Charbonneau, Janet Newman, Edward H. Snell, David R. So, Vincent Vanhoucke, Christopher J. Watkins, Shawn Williams, and Julie Wilson. Classification of crystallization outcomes using deep convolutional neural networks. PLOS ONE, 13(6): e0198883, June 2018. doi:10.1371/journal.pone.0198883.

25. D. Watts, K. Cowtan, and J. Wilson. Automated classification of crystallization experiments using wavelets and statistical texture characterization techniques. Journal of Applied Crystallography, 41(1):8–17, February 2008. doi:10.1107/S0021889807049308.

26. Jamie Milne, Chen Qian, David Hargreaves, Yinhai Wang, and Julie Wilson. Not getting in too deep: A practical deep learning approach to routine crystallisation image classification. PLOS ONE, 18(3):e0282562, 2023. doi:10.1371/journal.pone.0282562.

27. Keith B. Ward, Mary Ann Perozzo, and William M. Zuk. Automatic preparation of protein crystals using laboratory robotics and automated visual inspection. Journal of Crystal Growth, 90(1):325–339, July 1988. doi:10.1016/0022-0248(88)90328-4.

28. Paul V. C. Hough. Method and means for recognizing complex patterns, December 1962.

29. William M. Zuk and Keith B. Ward. Methods of analysis of protein crystal images. Journal of Crystal Growth, 110(1):148–155, March 1991. doi:10.1016/0022-0248(91)90878-9.

30. Christian A. Cumbaa, Angela Lauricella, Nancy Fehrman, Christina Veatch, Robert Collins, Joe Luft, George DeTitta, and Igor Jurisica. Automatic classification of sub-microlitre protein-crystallization trials in 1536-well plates. Acta Crystallographica Section D Biological Crystallography, 59(9):1619–1627, September 2003. doi:10.1107/S0907444903015130.

31. Teuvo Kohonen. Self-organized formation of topologically correct feature maps. Biological Cybernetics, 43(1):59–69, January 1982. doi:10.1007/BF00337288.

32. G. Spraggon, S. A. Lesley, A. Kreusch, and J. P. Priestle. Computational analysis of crystallization trials. Acta Crystallographica Section D: Biological Crystallography, 58(11):1915–1923, November 2002. doi:10.1107/S0907444902016840.

33. Roy Liu, Yoav Freund, and Glen Spraggon. Image-based crystal detection: A machine-learning approach. Acta Crystallographica Section D: Biological Crystallography, 64(Pt 12): 1187–1195, November 2008. doi:10.1107/S090744490802982X.

34. Christian A. Cumbaa and Igor Jurisica. Protein crystallization analysis on the World Community Grid. Journal of Structural and Functional Genomics, 11(1):61–69, March 2010. doi:10.1007/s10969-009-9076-9.

35. Jia Tsing Ng, Carien Dekker, Markus Kroemer, Michael Osborne, and Frank von Delft. Using textons to rank crystallization droplets by the likely presence of crystals. Acta Crystallographica Section D: Biological Crystallography, 70(Pt 10):2702–2718, September 2014. doi:10.1107/S1399004714017581.

36. Margot Yann and Yichuan Tang. Learning Deep Convolutional Neural Networks for X-Ray Protein Crystallization Image Analysis. Proceedings of the AAAI Conference on Artificial Intelligence, 30(1), February 2016. doi:10.1609/aaai.v30i1.10150.

37. E. H. Snell, J. R. Luft, S. A. Potter, A. M. Lauricella, S. M. Gulde, M. G. Malkowski, M. Koszelak-Rosenblum, M. I. Said, J. L. Smith, C. K. Veatch, R. J. Collins, G. Franks, M. Thayer, C. Cumbaa, I. Jurisica, and G. T. DeTitta. Establishing a training set through the visual analysis of crystallization trials. Part I: ∼150 000 images. Acta Crystallographica Section D: Biological Crystallography, 64(11):1123–1130, November 2008. doi:10.1107/S0907444908028047.

38. Soheil Ghafurian, Peter Orth, Corey Strickland, Hua Su, Sangita Patel, Steven Soisson, and Belma Dogdas. Classification of Protein Crystallization X-Ray Images Using Major Convolutional Neural Network Architectures, May 2018.

39. Kaiming He, Xiangyu Zhang, Shaoqing Ren, and Jian Sun. Deep Residual Learning for Image Recognition. In 2016 IEEE Conference on Computer Vision and Pattern Recognition (CVPR), pages 770–778, June 2016. doi:10.1109/CVPR.2016.90.

40. Formulatrix. Protein Crystallization Software Update: ROCK MAKER 3.15, July 2019.

41. Unversity at Buffalo. MARCO: MAchine Recognition of Crystallization Outcomes, 2018.

42. Yusei Miura, Tetsuya Sakurai, Claus Aranha, Toshiya Senda, Ryuichi Kato, and Yusuke Yamada. Classification of X-Ray Protein Crystallization Using Deep Convolutional Neural Networks with a Finder Module, December 2018.

43. Olaf Ronneberger, Philipp Fischer, and Thomas Brox. U-Net: Convolutional Networks for Biomedical Image Segmentation. In Nassir Navab, Joachim Hornegger, William M. Wells, and Alejandro F. Frangi, editors, Medical Image Computing and Computer-Assisted Intervention – MICCAI 2015, Lecture Notes in Computer Science, pages 234–241, Cham, 2015. Springer International Publishing. ISBN 978-3-319-24574-4. doi:10.1007/978-3-319-24574-4_28.

44. Olga Russakovsky, Jia Deng, Hao Su, Jonathan Krause, Sanjeev Satheesh, Sean Ma, Zhiheng Huang, Andrej Karpathy, Aditya Khosla, Michael Bernstein, Alexander C. Berg, and Li Fei-Fei. ImageNet Large Scale Visual Recognition Challenge. International Journal of Computer Vision, 115(3):211–252, December 2015. doi:10.1007/s11263-015-0816-y.

45. Mingxing Tan and Quoc V. Le. EfficientNet: Rethinking Model Scaling for Convolutional Neural Networks, September 2020.

46. David William Edwards II and Imren Dinc. Classification of Protein Crystallization Images using EfficientNet with Data Augmentation. In CSBio ‘20: Proceedings of the Eleventh International Conference on Computational Systems-Biology and Bioinformatics, CSBio2020, pages 54–60, New York, NY, USA, November 2020. Association for Computing Machinery. ISBN 978-1-4503-8823-8. doi:10.1145/3429210.3429220.

47. Nicholas Rosa, Christopher J. Watkins, and Janet Newman. Moving beyond MARCO. PLOS ONE, 18(3):e0283124, March 2023. doi:10.1371/journal.pone.0283124.

48. Francois Chollet. Xception: Deep Learning with Depthwise Separable Convolutions. In 2017 IEEE Conference on Computer Vision and Pattern Recognition (CVPR), pages 1800–1807, Honolulu, HI, July 2017. IEEE. ISBN 978-1-5386-0457-1. doi:10.1109/CVPR.2017.195.

49. Gao Huang, Zhuang Liu, Laurens Van Der Maaten, and Kilian Q. Weinberger. Densely Connected Convolutional Networks. In 2017 IEEE Conference on Computer Vision and Pattern Recognition (CVPR), pages 2261–2269. IEEE Computer Society, July 2017. ISBN 978-1-5386-0457-1. doi:10.1109/CVPR.2017.243.

50. Y. Thielmann, T. Luft, N. Zint, and J. Koepke. Crystal search – feasibility study of a real-time deep learning process for crystallization well images. Acta Crystallographica Section A: Foundations and Advances, 79(4):331–338, July 2023. doi:10.1107/S2053273323001948.

51. Alex Krizhevsky, Ilya Sutskever, and Geoffrey E Hinton. ImageNet Classification with Deep Convolutional Neural Networks. In Advances in Neural Information Processing Systems, volume 25. Curran Associates, Inc., 2012.

52. Karen Simonyan and Andrew Zisserman. Very Deep Convolutional Networks for Large-Scale Image Recognition, April 2015.

53. Forrest N. Iandola, Song Han, Matthew W. Moskewicz, Khalid Ashraf, William J. Dally, and Kurt Keutzer. SqueezeNet: AlexNet-level accuracy with 50x fewer parameters and <0.5MB model size, November 2016.

54. Marie-Noëlle Pons and Hervé Vivier. Crystallization monitoring by quantitative image analysis. Analytica Chimica Acta, 238:243–249, January 1990. doi:10.1016/S0003-2670(00)80543-7.

55. P. A. Larsen, J. B. Rawlings, and N. J. Ferrier. An algorithm for analyzing noisy, in situ images of high-aspect-ratio crystals to monitor particle size distribution. Chemical Engineering Science, 61(16):5236–5248, August 2006. doi:10.1016/j.ces.2006.03.035.

56. Zhenguo Gao, Yuanyi Wu, Ying Bao, Junbo Gong, Jingkang Wang, and Sohrab Rohani. Image Analysis for In-line Measurement of Multidimensional Size, Shape, and Polymorphic Transformation of l-Glutamic Acid Using Deep Learning-Based Image Segmentation and Classification. Crystal Growth & Design, 18(8):4275–4281, August 2018. doi:10.1021/acs.cgd.8b00883.

57. Kaiming He, Georgia Gkioxari, Piotr Dollár, and Ross Girshick. Mask R-CNN. IEEE Transactions on Pattern Analysis and Machine Intelligence, 42(2):386–397, February 2020. doi:10.1109/TPAMI.2018.2844175.

58. Daniel Bischoff, Brigitte Walla, and Dirk Weuster-Botz. Machine learning-based protein crystal detection for monitoring of crystallization processes enabled with large-scale synthetic data sets of photorealistic images. Analytical and Bioanalytical Chemistry, 414(21): 6379–6391, September 2022. doi:10.1007/s00216-022-04101-8.

59. Jiaming Han, Jian Ding, Jie Li, and Gui-Song Xia. Align Deep Features for Oriented Object Detection. IEEE Transactions on Geoscience and Remote Sensing, 60:1–11, 2022. doi:10.1109/TGRS.2021.3062048.

60. Jiangping Qin, Yan Zhang, Huan Zhou, Feng Yu, Bo Sun, and Qisheng Wang. Protein Crystal Instance Segmentation Based on Mask R-CNN. Crystals, 11(2):157, February 2021. doi:10.3390/cryst11020157.

61. Tsung-Yi Lin, Michael Maire, Serge Belongie, Lubomir Bourdev, Ross Girshick, James Hays, Pietro Perona, Deva Ramanan, C. Lawrence Zitnick, and Piotr Dollár. Microsoft COCO: Common Objects in Context, February 2015.

62. Kentaro Wada. Labelme: Image Polygonal Annotation with Python, November 2023.

63. Stephen M. Pizer, E. Philip Amburn, John D. Austin, Robert Cromartie, Ari Geselowitz, Trey Greer, Bart ter Haar Romeny, John B. Zimmerman, and Karel Zuiderveld. Adaptive histogram equalization and its variations. Computer Vision, Graphics, and Image Processing, 39(3):355–368, September 1987. doi:10.1016/S0734-189X(87)80186-X.

64. Zhuang Liu, Hanzi Mao, Chao-Yuan Wu, Christoph Feichtenhofer, Trevor Darrell, and Saining Xie. A ConvNet for the 2020s. In 2022 IEEE/CVF Conference on Computer Vision and Pattern Recognition (CVPR), pages 11966–11976. IEEE Computer Society, June 2022. ISBN 978-1-66546-946-3. doi:10.1109/CVPR52688.2022.01167.

65. Jeremy Howard and Sylvain Gugger. Fastai: A Layered API for Deep Learning. Information, 11(2):108, February 2020. doi:10.3390/info11020108.

66. Adam Paszke, Sam Gross, Francisco Massa, Adam Lerer, James Bradbury, Gregory Chanan, Trevor Killeen, Zeming Lin, Natalia Gimelshein, Luca Antiga, Alban Desmaison, Andreas Kopf, Edward Yang, Zachary DeVito, Martin Raison, Alykhan Tejani, Sasank Chilamkurthy, Benoit Steiner, Lu Fang, Junjie Bai, and Soumith Chintala. PyTorch: An Imperative Style, High-Performance Deep Learning Library. In H. Wallach, H. Larochelle, Beygelzimer, F. d\textbackslashtextquotesingle Alché-Buc, E. Fox, and R. Garnett, editors, Advances in Neural Information Processing Systems 32, pages 8024–8035. Curran Associates, Inc., 2019.

67. Ross Wightman. Pytorch image models. https://github.com/rwightman/pytorch-image-models, 2019.

68. Ilya Loshchilov and Frank Hutter. Decoupled Weight Decay Regularization, January 2019.

69. Leslie N. Smith and Nicholay Topin. Super-convergence: Very fast training of neural networks using large learning rates. In Artificial Intelligence and Machine Learning for Multi-Domain Operations Applications, volume 11006, pages 369–386. SPIE, May 2019. doi:10.1117/12.2520589.

70. Ramprasaath R. Selvaraju, Michael Cogswell, Abhishek Das, Ramakrishna Vedantam, Devi Parikh, and Dhruv Batra. Grad-CAM: Visual Explanations from Deep Networks via Gradient-Based Localization. In 2017 IEEE International Conference on Computer Vision (ICCV), pages 618–626, October 2017. doi:10.1109/ICCV.2017.74.

71. G. Bradski. The OpenCV library. Dr. Dobb’s Journal of Software Tools, 2000.

72. Robert Simpson, Kevin R. Page, and David De Roure. Zooniverse: Observing the world’s largest citizen science platform. In Proceedings of the 23rd International Conference on World Wide Web, WWW ‘14 Companion, pages 1049–1054, New York, NY, USA, April 2014. Association for Computing Machinery. ISBN 978-1-4503-2745-9. doi:10.1145/2567948.2579215.

73. Tilo Strutz. The Distance Transform and its Computation, February 2023.

74. Kaiming He, Xiangyu Zhang, Shaoqing Ren, and Jian Sun. Delving Deep into Rectifiers: Surpassing Human-Level Performance on ImageNet Classification. In 2015 IEEE International Conference on Computer Vision (ICCV), pages 1026–1034, Santiago, Chile, December 2015. IEEE. ISBN 978-1-4673-8391-2. doi:10.1109/ICCV.2015.123.

75. Gao Huang, Yu Sun, Zhuang Liu, Daniel Sedra, and Kilian Q. Weinberger. Deep networks with stochastic depth. In Bastian Leibe, Jiri Matas, Nicu Sebe, and Max Welling, editors, Computer Vision – ECCV 2016, pages 646–661. Springer International Publishing. ISBN 978-3-319-46493-0. doi:10.1007/978-3-319-46493-0_39.

